# Modular innate recognition in conventional T cells through FcRγ reconstitution

**DOI:** 10.64898/2026.11.20.768012

**Authors:** Caitlin Halligan, Suwen Li, Yee Nip, Jigar Patel, Jun Chan, Kevin Chen, William Harvey, Kristy Dai, Olivia Zhou, Joshua Hockman, Josephine C. K. Wong, Nikhil Sarvaiya, Vivek Chotai, Sanaz Memarzadeh, Caspian Oliai, Amélie Montel-Hagen, Sunmin Park, Jocelyn T. Kim, Christopher S. Seet

**Affiliations:** Department of Medicine, Division of Hematology-Oncology, David Geffen School of Medicine at UCLA, Los Angeles, California, USA; Department of Medicine, Division of Infectious Diseases, David Geffen School of Medicine at UCLA, Los Angeles, California, USA; Department of Obstetrics and Gynecology, David Geffen School of Medicine at UCLA, Los Angeles, California, USA; VA Greater Los Angeles Healthcare System, Los Angeles, CA, 90073, USA; Department of Molecular and Medical Pharmacology, David Geffen School of Medicine at UCLA, Los Angeles, California, USA; Jonsson Comprehensive Cancer Center, David Geffen School of Medicine at UCLA, Los Angeles, California, USA; Broad Stem Cell Research Center, David Geffen School of Medicine at UCLA, Los Angeles, California, USA; Molecular Biology Institute, University of California Los Angeles, California, USA

**Author notes:** Department of Hematology & Hematopoietic Cell Transplantation, City of Hope, Duarte, California, USA. These authors contributed equally to this work (Caitlin Halligan, Suwen Li).

## Abstract

Engineered T cells can elicit durable tumor responses but largely rely on TCRs or CARs for target recognition. Here we show that FcRγ, an innate signaling adaptor not expressed by conventional T cells, enables full-length innate receptor function in adult human αβ T cells. FcRγ reconstitution enabled endogenous NKp30 surface expression and cytotoxicity without inducing an innate-like state. Stable co-expression of NKp30 and FcRγ maintained receptor expression after activation withdrawal, supported sustained tumor killing, delayed leukemia progression in mice, and enabled cytotoxicity against primary myeloid leukemia. FcRγ also supported functional import of receptors spanning natural cytotoxicity, Fc receptor, and C-type lectin families, extending conventional T cell responses to tumor ligands, antibody-coated targets, IgE receptor engagement, and microbial patterns. These findings establish a shared-adaptor strategy for expanding innate recognition in engineered T cells while preserving the identity and functional properties of the widely adopted conventional T cell platform.

## Introduction

Conventional αβ T cells are the most clinically validated platform for engineered cellular immunotherapy. Their capacity for expansion, tissue trafficking, and long-term persistence has enabled durable responses and established standard-of-care therapies in several advanced malignancies^1,2^. Target recognition, however, relies on engineered TCRs, which require intact peptide-MHC presentation, or CARs, which recognize predefined cell-surface antigens. TCR-based therapies are thus susceptible to defects in antigen processing and presentation, while both approaches remain vulnerable to heterogeneous target expression and antigen loss. These limitations underscore a need to identify additional routes of target recognition in conventional T cells.

NK cells, innate-like T cells (including γδ T cells, iNKT cells, and intraepithelial lymphocytes), and subsets of myeloid cells use diverse innate receptors to detect ligands associated with cellular stress, antibody-coated targets, and conserved microbial structures^3,4^. These broad recognition modes have motivated their development as alternative cellular therapies^5,6^. Many activating innate receptors lack autonomous signaling domains and instead depend on ITAM-containing adaptors, including the Fc receptor common γ-chain (FcRγ, encoded by FCER1G), CD3ζ, and DAP12, for receptor assembly and signal transduction^4,7–10^. Conventional αβ T cells use homologous ITAM-based signaling machinery downstream of TCR/CD3ζ activation but normally lack FcRγ and DAP12. Previous studies have introduced individual innate receptors or receptor-derived constructs into T cells^11–14^, whereas endogenous FcRγ and FcRγ-associated receptor expression have been described in distinct innate-like CD8+ αβ T cell populations^15,16^. We therefore asked whether FcRγ reconstitution could broadly enable innate receptor function in conventional T cells without requiring or inducing an innate-like state.

Here, we show that FcRγ reconstitution expands innate recognition in conventional adult human T cells while maintaining conventional T cell identity and function. Unexpectedly, we found that both CD8⁺ and CD4+ T cells upregulate NKp30, an NK-associated natural cytotoxic receptor (NCR), following activation but retain it intracellularly. FcRγ reconstitution induced NKp30 surface localization and enabled competent, ligand-dependent tumor cytotoxicity without a detectable shift toward an innate-like T cell state (Fig. 1a). Stable co-expression of NKp30 and FcRγ maintained receptor expression after activation withdrawal and supported sustained antitumor activity, outperforming primary NK cells under similar conditions. FcRγ also supported the functional expression of full-length innate receptors spanning natural cytotoxicity, Fc receptor, and C-type lectin families (Fig. 1b), enabling tumor, antibody, and microbial pattern recognition in T cells. Together, these findings highlight the potential for new targeting capabilities of innate receptors when deployed outside their native regulatory contexts, and establish a shared-adaptor engineering approach for expanding innate recognition in conventional T cells.

**Fig 1:**
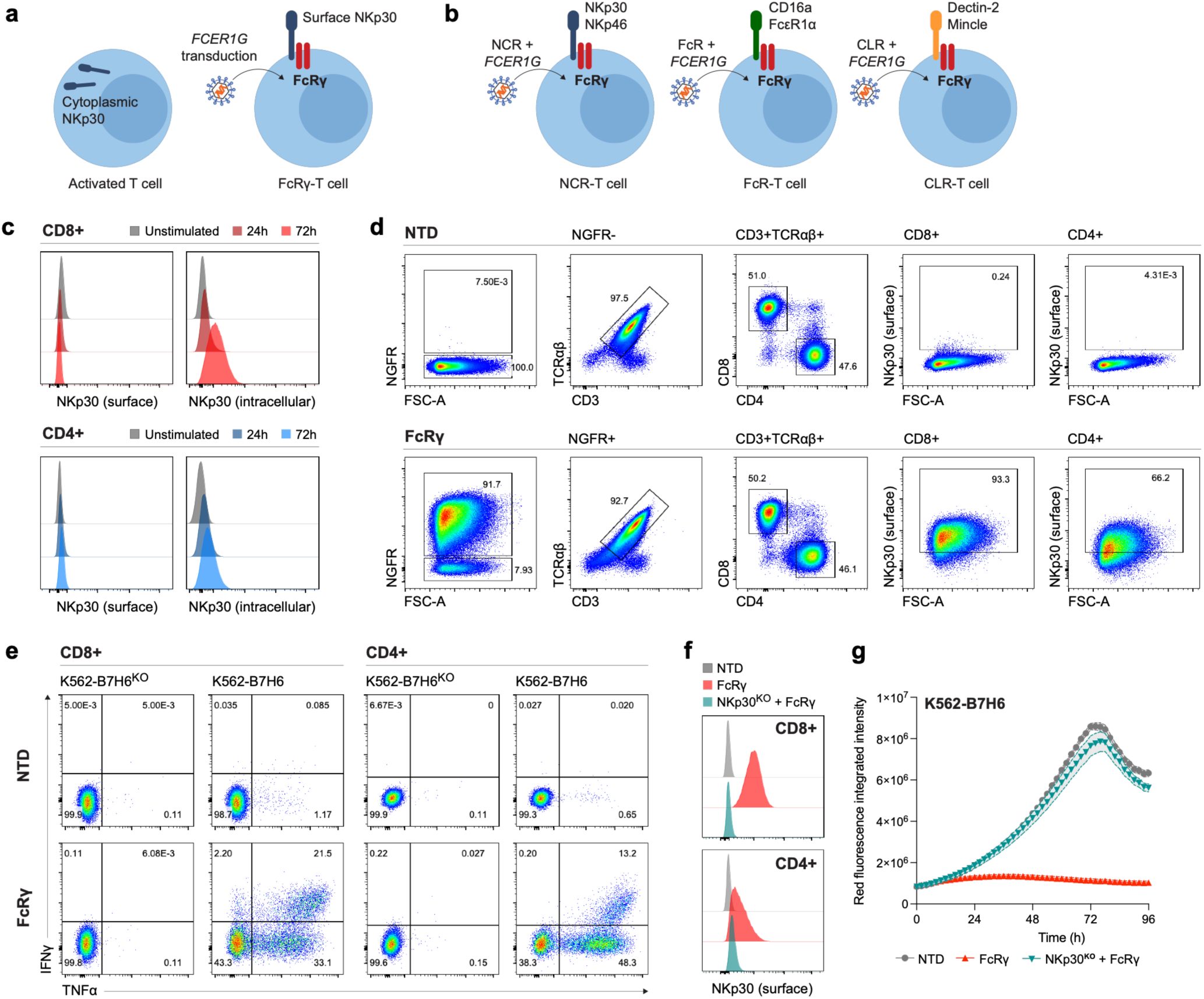
FcRγ enables functional endogenous NKp30 surface expression in activated conventional T cells. **a)** Graphical summary: FcRγ induces surface expression of endogenous NKp30 in activated T cells. **b)** Graphical summary: FcRγ enables functional expression of multiple classes of heterologous immune receptors in T cells. **c)** Flow cytometry of surface or intracellular NKp30 expression in CD8+ and CD4+ T cells. T cells were activated with anti-CD3/28 beads and IL-2 for the indicated times. Relative frequencies shown are normalized to mode. **d)** Flow cytometry expression of surface NKp30 and phenotype of non-transduced (NTD) or FcRγ -transduced T cells. NGFR is used as a marker of transduction. **e)** Intracellular IFNγ and TNFα expression in CD8+ and CD4+ NTD or FcRγ-transduced T cells after co-culture with B7-H6-knockout (K562-B7H6^KO^) or B7-H6-overexpressing (K562-B7H6) cell lines for 6 hours. **f)** Surface expression of NKp30 on NTD, FcRγ, or FcRγ-NKp30-knockout (NKp30^KO^) T cells. Relative frequencies shown are normalized to mode. **g)** IncuCyte tracking of mKate-labeled K562-B7H6 cells co-cultured with NTD, FcRγ, or FcRγ-NKp30^KO^ total T cells at 1:1 effector-to-target (E:T) ratio. Red fluorescence integrated intensity is shown. Data are mean ± SEM of triplicate wells.

## Results

### FcRγ induces NKp30 surface localization and function in activated T cells

In profiling activated, conventional αβ T cells from healthy adult peripheral blood donors, we unexpectedly detected broad intracellular expression of NKp30, an activating NCR associated with NK and innate-like T cells. NKp30 was induced in both CD8^+^ and CD4^+^ T cells following stimulation with anti-CD3/CD28 in the presence of either IL-2 (Fig. 1c) or IL-7/IL-15 (Fig. S1a), but was not detected at the cell surface, suggesting that NKp30 expression alone is insufficient for functional receptor assembly in conventional T cells.

NKp30 associates with CD3ζ and/or FcRγ dimers to form functional receptor-adaptor complexes^4,9^. Conventional T cells express CD3ζ but not FcRγ, suggesting that FcRγ availability might limit surface localization of intracellular NKp30. Supporting this possibility, FcRγ was previously shown to enable NKp30 surface expression in a subset of IL-15-cultured CD8⁺ T cells, although the function and identity of those cells following introduction of FcRγ were not reported^16^. We therefore tested whether FcRγ alone could enable endogenous NKp30 function in activated conventional T cells. Following lentiviral expression in activated, healthy donor T cells, FcRγ induced robust NKp30 surface localization in both CD8⁺ and CD4⁺ T cell subsets (Fig. 1d). Surface TCRαβ expression was minimally affected, suggesting that surface NKp30-FcRγ expression did not substantially interfere with TCR assembly, for example by competing for endogenous CD3ζ.

We next determined whether cell surface-localized NKp30 on FcRγ-T cells mediates target recognition and effector function. In co-culture experiments, both CD8⁺ and CD4⁺ FcRγ-T cells mounted ligand-specific cytokine responses to K562 cells engineered to overexpress the NKp30 ligand B7-H6 (K562-B7H6), but not to K562 cells in which B7-H6 was knocked out (K562-B7H6^KO^) (Fig. 1e; Fig. S1b). FcRγ-T cells also mediated potent cytotoxicity against K562-B7H6 cells, which was abrogated by CRISPR deletion of *NCR3*, the gene encoding NKp30, confirming NKp30-dependent killing (Fig. 1f,g).

Because innate-like T cell subtypes can express FcRγ^15,16^, we examined whether FcRγ reconstitution induced or selected for a broader innate-like T cell state. In activated T cells, FcRγ selectively induced surface expression of NKp30 but not other FcRγ-associated innate receptors, including NKp46, CD16a, CD64, and FcεRIα (Fig. S1c). Upregulation of the NK/innate-like T cell marker CD56 was not observed (Fig. S1c). Single-cell RNA sequencing of T cells from two healthy donors showed similar transcriptional clusters between non-transduced and FcRγ-T cells (Fig. S1d), with preserved expression of canonical T cell identity genes, including *CD3D* and *TRAC* (Fig. S1e). Expression of *TCF7* (encoding TCF1), associated with conventional naïve and memory T cells^15,17,18^, and *ZBTB16* (encoding PLZF), associated with innate-like lymphocyte differentiation^19–21^, was also unaltered (Fig. S1e). Consistent with flow cytometry, *NCR3*, encoding NKp30, was highly expressed in activated T cells independently of FcRγ transduction, whereas *NCR1* and *NCR2*, encoding NKp46 and NKp44, respectively, were not detected in either condition (Fig. S1e). Indeed, pseudobulk analysis identified *FCER1G* as the only significantly differentially expressed gene between FcRγ-T and non-transduced T cells (Fig. S1f). Together, these data demonstrate that FcRγ reconstitution enables endogenous NKp30-dependent cytotoxicity in activated conventional T cells without eliciting an innate-like T cell state.

### NKp30-FcRγ engineering creates stable, ligand-gated NCR-T cells

Although FcRγ enabled functional surface localization of endogenous NKp30 in activated T cells, NKp30 surface expression remained activation dependent, declining markedly following withdrawal of cytokines and TCR stimulation *in vitro* (Fig. 2a) and *in vivo* following adoptive transfer into NSG mice (Fig. S2a). To establish stable NKp30 activity in T cells, we therefore co-expressed NKp30 and FcRγ from a single lentiviral vector. These NKp30-FcRγ “NCR-T cells” exhibited higher surface NKp30 levels than FcRγ-T cells and, importantly, maintained NKp30 expression in the absence of exogenous cytokines or TCR stimulation (Fig. 2a; Fig. S2b). Using K562-B7H6 and K562-B7H6^KO^ target cells, we confirmed that NCR-T cells mounted B7-H6-specific cytokine responses (Fig. 2b) and cytotoxicity (Fig. 2c) in the absence of exogenous cytokines.

**Fig 2:**
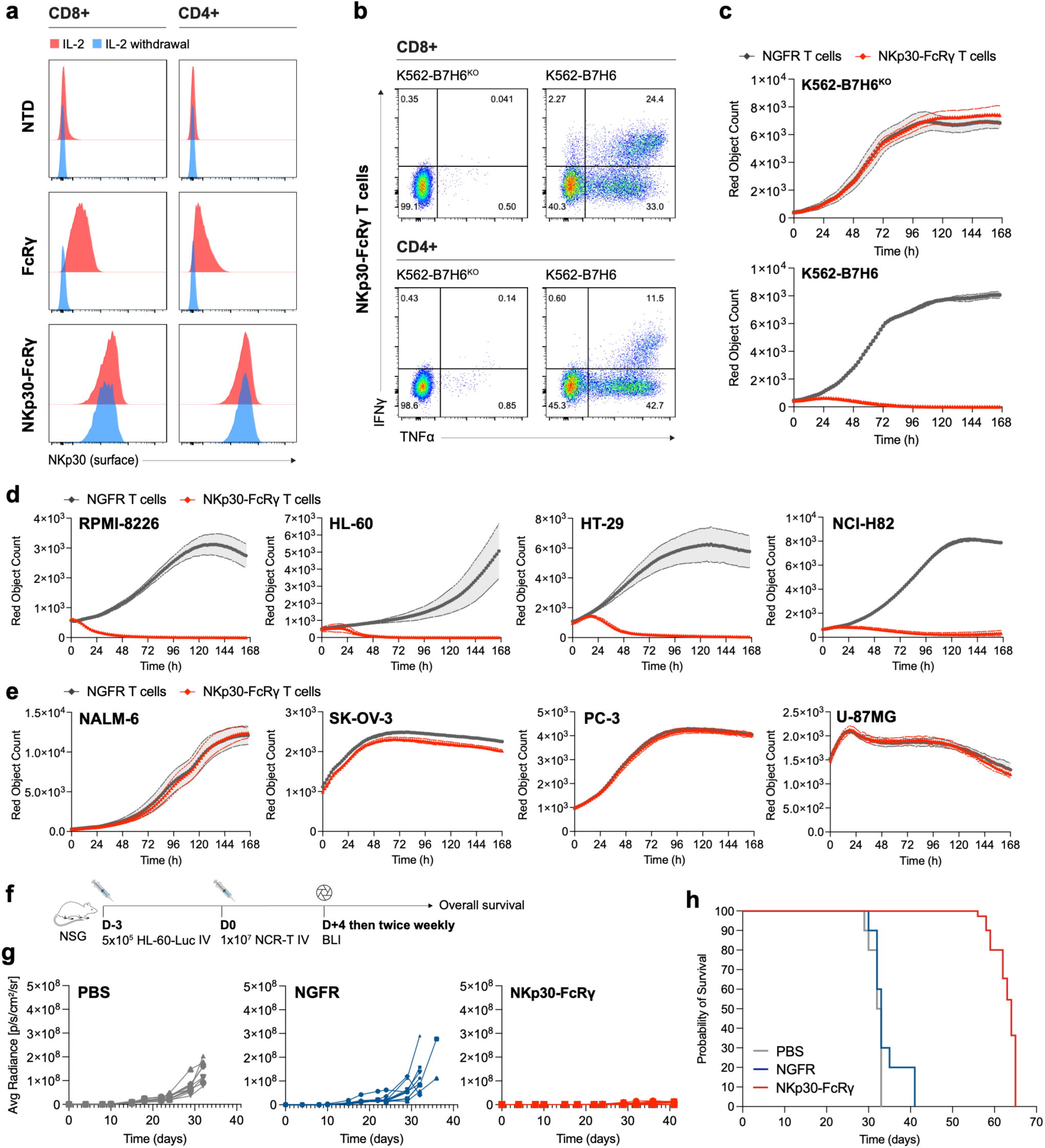
Stable NKp30-FcRγ engineering creates NCR-T cells with anti-tumor activity. **a)** Surface NKp30 expression in CD8+ and CD4+ T cells transduced with FcRγ alone or NKp30 and FcRγ (NKp30-FcRγ). Relative frequencies shown are normalized to mode. **b)** Intracellular IFNγ and TNFα expression in CD8+ and CD4+ NKp30-FcRγ NCR-T cells after 6h co-culture with B7-H6-knockout (K562-B7H6^KO^) or B7-H6-overexpressing (K562-B7H6) cell lines. **c)** IncuCyte counts of mKate-labeled K562-B7H6^KO^ or K562-B7H6 cells co-cultured with NGFR-only or NKp30-FcRγ transduced total T cells at a 1:1 E:T without exogenous cytokines. Data are mean ± SEM of triplicate wells. **d)** IncuCyte counts of mKate-transduced tumor cell lines (described in Supplementary Data 1) co-cultured with *TRAC*^KO^ NGFR-only or *TRAC*^KO^ NKp30-FcRγ total NCR-T cells at a 1:1 E:T without exogenous cytokines. Representative cell lines killed by NKp30-FcRγ NCR-T cells are shown. Data are mean ± SEM of triplicate wells. **e)** IncuCyte assays, as described in panel d, for representative cell lines not killed by NKp30-FcRγ NCR-T cells. Data are mean ± SEM of triplicate wells. **f)** Schematic of HL-60 AML xenograft/adoptive cell therapy mouse model. **g)** Tumor responses by luciferase bioluminescence imaging (BLI) in HL-60-luciferase engrafted NSG mice, as described in panel f. Mice received PBS (no effector cell control), total NGFR-only T cells, or total NKp30-FcRγ NCR-T cells (*n*=10 per group). **h)** Kaplan-Meier survival probability curves of mice shown in panel g.

As B7-H6 is expressed across human malignancies^22,23^, we evaluated the cytotoxic activity of NCR-T cells against a large panel of cancer cell lines in the absence of exogenous cytokines. NCR-T cells effected broad cytotoxicity across hematologic and solid tumor types, including multiple myeloma, AML, T-ALL, B cell lymphoma, small cell lung cancer, ovarian, cervical, and colorectal cancer cell lines (Fig. 2d; Fig. S3a; Supplementary Data 1). Susceptibility was not universal, however, with resistance more frequently observed among the B-ALL, prostate cancer, glioma, and breast cancer lines tested (Fig. 2e; Fig. S3b). In at least one resistant cell line, M202 melanoma, B7-H6 overexpression restored susceptibility to NCR-T cell killing, suggesting that ligand abundance can be a determinant of response (Fig. S3c,d,f).

Correlating NCR-T cell susceptibility with tumor cell B7-H6 expression proved challenging, as endogenous B7-H6 was poorly detected on tumor cell lines by flow cytometry (Fig. S3e). B7-H6 abundance measured by Western blotting generally tracked with susceptibility but did not fully account for response heterogeneity (Fig. S3g). Furthermore, higher-molecular-weight B7-H6 species were observed and varied across cell lines, potentially reflecting post-translational modifications, such as glycosylation^24^; however, the specific contributions of ligand abundance and post-translational processing to NCR-T cell recognition remain to be determined.

In an HL-60 AML mouse xenograft model, adoptively transferred NCR-T cells significantly reduced leukemia progression and prolonged overall survival (Fig. 2g,h). Although NCR-T cell recipient mice ultimately succumbed to progressive disease, NCR-T cells remained detectable in the spleen, blood, and bone marrow at relapse and retained NKp30 surface expression (Fig. S4a,b). Disease progression therefore occurred despite persistent NCR-T cells and surface NKp30, although the mechanisms driving resistance remain to be determined. Together, these data establish NKp30-FcRγ as a stable receptor-adaptor module in conventional T cells for innate tumor recognition.

### NKp30-FcRγ NCR-T cells show sustained activity compared with primary NK cells

Primary NK cells natively express NKp30 but often exhibit limited *in vivo* persistence without cytokine support^25,26^. We hypothesized that NKp30-FcRγ NCR-T cells would combine innate tumor recognition with the persistence of conventional T cells, particularly under cytokine-limited conditions. We therefore compared the *in vitro* and *in vivo* durability of antitumor responses of NCR-T cells and donor-matched expanded primary NK cells in the absence of exogenous cytokines. At baseline, NCR-T cells expressed higher levels of surface NKp30 than expanded NK cells (Fig. 3a; Fig. S5a). In serial killing assays against the RPMI-8226 myeloma cell line, which expresses B7-H6 and is a target of NCR-T cell killing (Fig. S3e,g), NCR-T cell cytotoxicity was maintained across repeated tumor challenges, whereas NK cell cytotoxicity was rapidly lost following the first tumor challenge (Fig. 3b). In a stringent HL-60 AML xenograft model, primary NK cells conferred no improvement in tumor control or survival over PBS-treated controls, whereas NCR-T cells significantly reduced tumor progression and prolonged survival (Fig. 3c,d). Endpoint analysis revealed undetectable NK cells but persistent NCR-T cells in the blood and spleen of respective recipient mice (Fig. S5b,c). Together, these findings demonstrate that NCR-T cells sustain antitumor activity and persist longer *in vivo* than expanded primary NK cells under the conditions tested.

**Fig 3:**
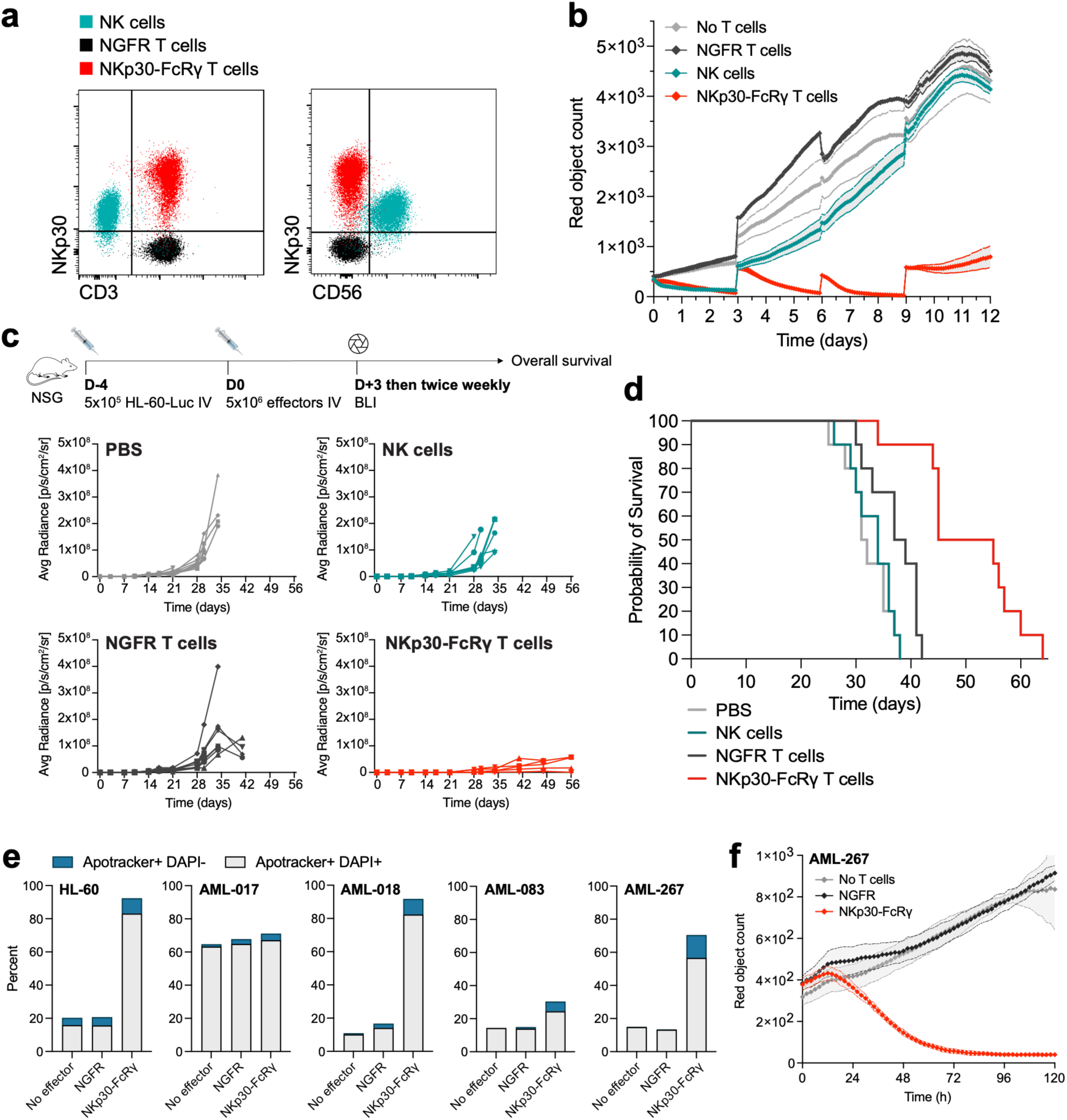
NKp30-FcRγ NCR-T cells show sustained activity compared with primary NK cells. **a)** Flow cytometry overlays showing expression of surface NKp30 on primary NK cells expanded with K562-mbIL21 cells and IL-2/IL-15, and same-donor T cells transduced with NGFR-only or NKp30-FcRγ. Expression of T cell (CD3) and NK cell (CD56) markers are shown. **b)** IncuCyte counts of mKate-transduced RPMI-8226 myeloma cells co-cultured with primary NK cells or same-donor total T cells transduced with NGFR-only or NKp30-FcRγ, as described in panel a, at a 1:1 E:T without exogenous cytokines. The starting number of tumor cells was added to wells every 72 h to assess serial killing capacity. Data are mean ± SEM of triplicate wells. **c)** Schematic of HL-60 AML xenograft/adoptive cell therapy NSG mouse model, and tumor response plots of luciferase bioluminescence imaging of HL-60-luciferase engrafted mice which received PBS (no effector control), primary NK cells, or same-donor total T cells transduced with NGFR or NKp30-FcRγ, as described in panel a. **d)** Kaplan-Meier survival probability curves of mouse cohorts shown in panel c. **e)** Apoptosis assays of HL-60 AML cells (positive control) and primary CD34+ AML blasts showing percent of early (apotracker+ DAPI-) and late (apotracker+ DAPI+) apoptotic cells after co-culture with healthy donor *TRAC*^KO^ NGFR-only or NKp30-FcRγ total NCR-T cells at a 1:1 E:T for 72 hours. **f)** IncuCyte counts of mKate-transduced primary AML blasts from a single patient, co-cultured with healthy donor *TRAC*^KO^ NGFR-only or NKp30-FcRγ total NCR-T cells at a 1:1 E:T. Data are mean ± SEM of triplicate wells.

### NKp30-FcRγ NCR-T cells target primary myeloid leukemia

We next evaluated cytotoxicity against primary AML, where engineered T cell therapies face challenges from antigen heterogeneity and on-target, off-tumor toxicity^27,28^. CD34⁺ leukemic blasts were isolated from three patients with AML and one with blast-phase myeloproliferative neoplasm (Supplementary Table 1). NKp30-FcRγ NCR-T cells were generated from healthy donor T cells with additional CRISPR-mediated deletion of *TRAC* to reduce alloreactivity against patient samples (Fig. S5d). *In vitro*, *TRAC*-disrupted NCR-T cells mediated cytotoxicity against both HL-60 (control) and primary leukemia cells, whereas vector control T cells showed no specific killing (Fig. 3e,f; Fig. S5e). Of note, targeted leukemias included those with adverse-risk features, including *TP53*-mutated AML with complex cytogenetics, and *JAK2*-mutated blast-phase MPN. The magnitude of cytotoxicity varied across specimens but generally tracked with B7-H6 abundance by Western blot (Fig. 3e,f; Fig. S5f). Together, these findings extend NCR-T cytotoxicity to primary myeloid leukemia, including molecularly adverse risk disease.

### NKp30-FcRγ engagement couples innate recognition to TCR-like signaling outputs

FcRγ signaling in innate cells and TCR signaling in T cells rely on distinct but homologous ITAM-based signaling components^10,29,30^. We therefore hypothesized that NKp30-FcRγ could engage endogenous T cell signaling machinery to activate canonical TCR-like transcriptional outputs. In NCR-T cells transduced with reporters of NFAT, NF-κB, or AP-1 activity, B7-H6 engagement activated all three transcriptional reporters, closely mimicking the responses to cognate peptide-MHC engagement of TCR-T cells expressing 1G4, a high-affinity HLA-A*02:01/NY-ESO-1-specific TCR^31^ (Fig. S6a).

Given the importance of antigen receptor signals in determining T cell persistence and exhaustion, we compared the effects of chronic NKp30-FcRγ and TCR stimulation on exhaustion-associated phenotypes and cytokine-production capacity. NCR-T cells were subjected to repeated stimulation with K562-B7H6 cells in the absence of exogenous cytokines; in parallel, donor-matched 1G4 TCR-T cells underwent the same stimulation regimen using K562-ESO SCT cells. After four serial 3-day stimulations, NCR-T cells exhibited lower expression of several exhaustion-associated markers and reduced marker co-expression than 1G4 TCR-T cells (Fig. S6b–d). NCR-T cells also retained a higher frequency of polyfunctional cells producing combinations of IFNγ, TNFα, and IL-2 in response to PMA/ionomycin (Fig. S6e,f). Thus, NKp30-FcRγ engagement activates canonical T cell signaling outputs in NCR-T cells while maintaining their functional capacity in models of chronic antigen stimulation.

### NKp46-FcRγ extends tumor recognition and reveals context-dependent reactivity

We next tested whether FcRγ reconstitution could enable the functional import of additional innate receptors into T cells. NKp46 is an NCR expressed by NK cells and certain innate-like T cells, where it associates with dimeric FcRγ and/or CD3ζ signaling adaptors^4,32^. We confirmed that NKp46 mRNA was not significantly expressed in conventional T cells following activation (Fig. S1e), nor was surface expression induced following FcRγ transduction (Fig. S1c). NKp46 transduction alone resulted in minimal surface expression, whereas co-transduction with FcRγ markedly increased NKp46 surface localization in both CD8⁺ and CD4⁺ T cells while also preserving surface TCR levels (Fig. 4a,b; Fig. S7a). Concurrent surface NKp30 expression was also observed, consistent with FcRγ-induced membrane localization of endogenous NKp30.

**Fig 4:**
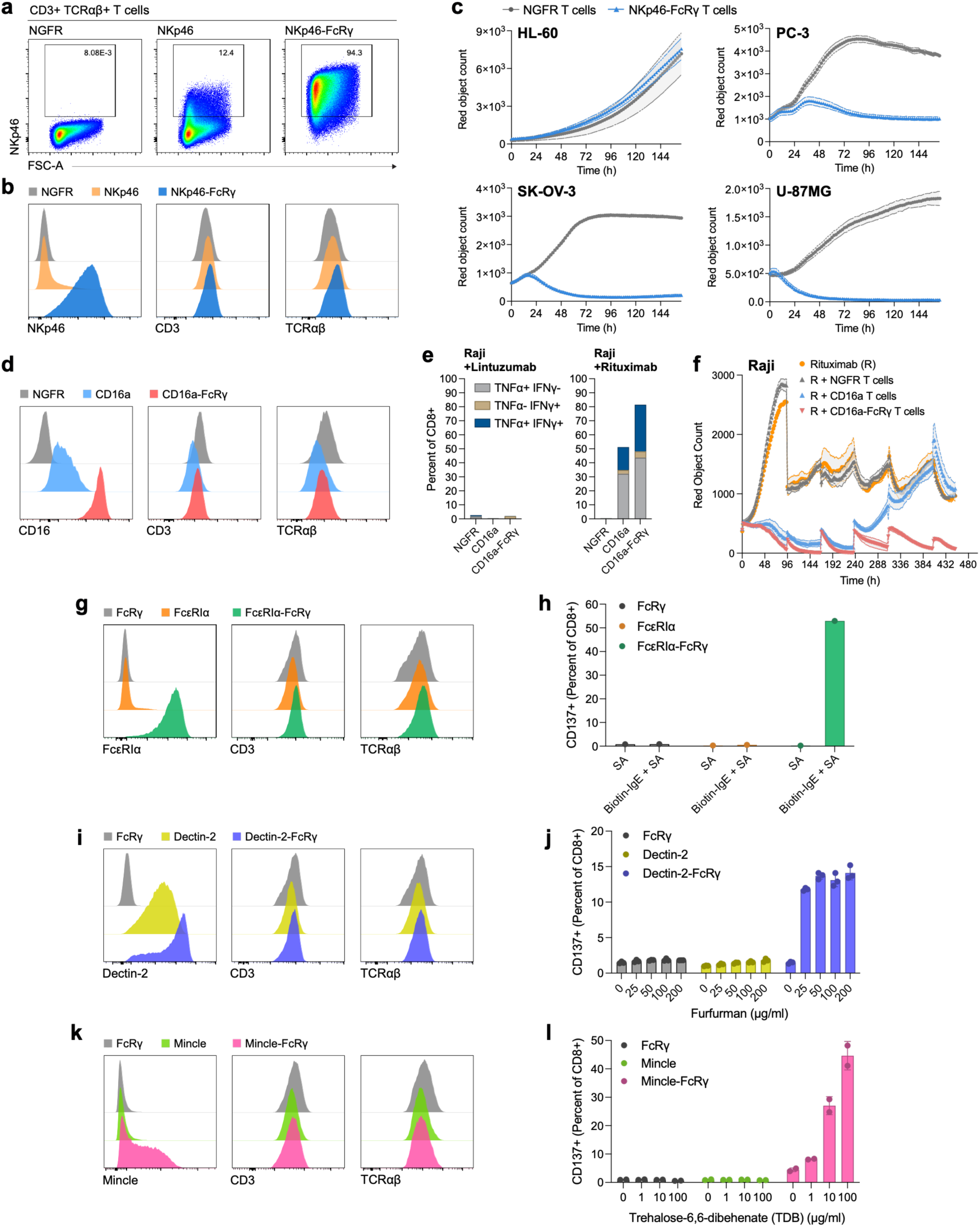
FcRγ reconstitution enables the functional import of heterologous immune receptors into conventional T cells. **a)** Flow cytometry expression of surface NKp46 on total T cells transduced with NGFR-only, NKp46 alone, or NKp46 with FcRγ (NKp46-FcRγ). **b)** Expression levels of NKp46, CD3, and TCRαβ on NGFR, NKp46, or NKp46-FcRγ total T cells described in panel a. Relative frequencies shown are normalized to mode. **c)** IncuCyte counts of mKate-transduced tumor cell lines co-cultured with *TRAC*^KO^ NGFR-only or *TRAC*^KO^ NKp46-FcRγ total NCR-T cells at a 1:1 E:T without exogenous cytokines. Data are mean ± SEM of triplicate wells. **d)** Expression levels of CD16a, CD3, and TCRαβ on total T cells transduced with NGFR, CD16a alone, or CD16a with FcRγ (CD16a-FcRγ). Relative frequencies shown are normalized to mode. **e)** Intracellular IFNγ and TNFα expression in CD8+ T cells transduced with NGFR, CD16a, or CD16a-FcRγ after co-culture with Raji cells for 6 h in the presence of lintuzumab (anti-CD33) or rituximab (anti-CD20) antibodies. **f)** IncuCyte counts of mKate-transduced Raji cells co-cultured with total T cells transduced with NGFR, CD16a, or CD16a-FcRγ at a 1:1 E:T without exogenous cytokines in the presence of Rituximab (R). The starting number of Raji cells was added to the cultures every 72 h to assess serial killing capacity. Data are mean ± SEM of triplicate wells. **g)** Expression levels of FcεRIα, CD3, and TCRαβ on total T cells transduced with FcRγ alone, FcεRIα, or FcεRIα with FcRγ (FcεRIα-FcRγ). Relative frequencies shown are normalized to mode. **h)** Activation-induced CD137 expression on CD8+ T cells transduced with FcRγ alone, FcεRIα, or FcεRIα with FcRγ cultured for 24h on streptavidin-only or streptavidin + biotin-IgE-Fc-coated plates. **i)** Expression levels of Dectin-2, CD3, and TCRαβ on total T cells transduced with FcRγ alone, Dectin-2, or Dectin-2 with FcRγ (Dectin-2-FcRγ). Relative frequencies shown are normalized to mode. **j)** Activation-induced CD137 expression on CD8+ T cells transduced with FcRγ alone, Dectin-2, or Dectin-2-FcRγ cultured for 18 h with Furfurman at the indicated concentrations. **k)** Expression levels of Mincle (CLEC4E), CD3, and TCRαβ on T cells transduced with FcRγ alone, Mincle, or Mincle with FcRγ (Mincle-FcRγ). Relative frequencies shown are normalized to mode. **l)** Activation-induced CD137 expression on CD8+ T cells transduced with FcRγ alone, Mincle, or Mincle-FcRγ cultured for 18 h with trehalose-6,6-dibehenate (TDB) at the indicated concentrations.

The ligands responsible for human NKp46 recognition remain incompletely defined, complicating assessment of ligand-specific function in NKp46-FcRγ T cells. Cell surface-exposed calreticulin (ectoCRT) has been reported as a candidate NKp46 ligand^33^. In co-culture experiments, NKp46-FcRγ T cells effected cytotoxicity against K562-B7H6 cells, consistent with surface expression of endogenous NKp30, and not against K562-B7H6^KO^ cells. Expression of human GPI-anchored ectoCRT in K562-B7H6^KO^ cells, however, did not confer susceptibility under the conditions tested (Fig. S7b,c). Nevertheless, NKp46-FcRγ T cells mediated robust cytotoxicity against multiple tumor cell lines, including epithelial and glioma lines resistant to NKp30-FcRγ NCR-T cell killing (Fig. 4c; Fig. S7d). NKp46-FcRγ T cells and NKp30-FcRγ NCR-T cells displayed distinct patterns of tumor cell targeting: in cytokine-free killing assays, HL-60 cells were susceptible to NKp30-FcRγ but resistant to NKp46-FcRγ T cells, whereas NKp30-FcRγ-resistant cell lines including PC-3, SK-OV-3, OVCAR-8, and U-87MG showed the opposite susceptibility pattern (Fig. 4c; Fig. S7d).

*In vivo*, NKp46-FcRγ T cells induced an initial, modest antitumor response in a NALM-6 B-ALL mouse xenograft model (Fig. S7f) but were associated with early morbidity and mortality compared with non-transduced T cells (Fig. S7g). Necropsy revealed robust splenic persistence of human T cells accompanied by dense T cell infiltrates in the liver and lungs (Fig. S7h); however, the mechanism underlying these findings remains to be determined. Together, these results demonstrate that FcRγ-enabled NKp46 expression expands solid tumor recognition in conventional T cells while highlighting the need to define receptor-specific targeting and tissue reactivity when expressed in the context of conventional T cells.

### FcRγ reconstitution improves antibody-dependent cytotoxicity in T cells through CD16a

We next asked whether FcRγ adaptor reconstitution extends beyond NCRs to support Fc receptor (FcR) function in T cells. CD16a (FcγRIIIa), an IgG receptor, mediates antibody-dependent cellular cytotoxicity (ADCC) in NK cells through association with FcRγ and/or CD3ζ^7,30,34^. CD16a alone can be ectopically expressed and support ADCC in human T cells, with a previous report showing co-expression of either FcRγ or CD3ζ increased receptor surface expression, with functional effects varying by antibody-target combination^14^. In our system, CD16a transduction alone yielded modest surface expression with reduced TCRαβ and CD3ε levels, suggesting competition with TCR assembly for endogenous CD3ζ (Fig. 4d; Fig. S8a). Co-expression with FcRγ markedly enhanced CD16a surface expression while preserving TCRαβ and CD3ε levels (Fig. 4d; Fig. S8a). Compared with CD16a alone, CD16a-FcRγ “FcR-T” cells mounted enhanced cytokine responses to both soluble and plate-bound CD16 agonist antibody (Fig. S8b), as well as to the CD20-high Raji and Ramos lymphoma cell lines in the presence of the anti-CD20 antibody rituximab, but not the irrelevant anti-CD33 antibody lintuzumab (Fig. 4e; Fig. S8c,d). FcRγ co-expression notably enabled T cell activation and ADCC against the CD20-low NALM-6 B-ALL cell line, which was poorly recognized by T cells expressing CD16a alone (Fig. S8c,e,f). CD16a-FcRγ FcR-T cells also exhibited more sustained ADCC across serial challenges with Raji cells (Fig. 4f) and greater proliferation in response to rituximab-coated target cells, including NALM-6, compared with CD16a alone (Fig. S8g). Together, these data show that FcRγ enhances CD16a surface expression, and CD16-mediated activation, sustained ADCC, and antibody-dependent T cell expansion in conventional T cells.

### FcRγ enables high-affinity IgE receptor function in conventional T cells

We also tested whether FcRγ reconstitution enables functional T cell expression of FcεRIα, the IgE-binding subunit of the high-affinity IgE receptor^35^. In mast cells and basophils, FcεRIα associates with FcεRIβ and FcRγ, although the β chain is dispensable for surface expression of the human receptor^35,36^. When transduced alone into T cells, FcεRIα showed minimal surface expression. Co-transduction with FcRγ, however, resulted in robust FcεRIα surface expression on both CD8⁺ and CD4⁺ T cells without altering TCRαβ or CD3ε levels (Fig. 4g; Fig. S9a). To assess receptor function, we stimulated T cells with biotinylated human IgE-Fc immobilized on streptavidin-coated plates^37^. Immobilized IgE-Fc induced upregulation of the T cell activation marker CD137 on both CD8⁺ and CD4⁺ FcεRIα-FcRγ T cells, whereas T cells transduced with FcεRIα alone showed no detectable response (Fig. 4h; Fig. S9b). These data demonstrate that FcRγ enables FcεRIα surface expression and IgE-dependent activation in T cells.

### FcRγ extends T cell recognition to microbial molecular patterns

Finally, we tested whether FcRγ supports the function of myeloid pattern-recognition receptors in conventional T cells. Dectin-2 (*CLEC6A*), a C-type lectin receptor (CLR), recognizes mannose-containing cell wall components of fungi and mycobacteria. In myeloid cells, Dectin-2 associates with FcRγ to drive pro-inflammatory cytokine responses and antifungal immunity^38–41^. When transduced alone into T cells, Dectin-2 exhibited moderate surface expression (Fig. 4i; Fig. S9c) but failed to induce T cell activation in response to furfurman, a cell wall preparation from *Malassezia furfur* (Fig. 4j; Fig. S9d). Co-expression of FcRγ both markedly increased Dectin-2 surface expression on CD8⁺ and CD4⁺ T cells (Fig. 4i; Fig. S9c) and enabled furfurman-induced activation, as measured by CD137 upregulation (Fig. 4j; Fig. S9d).

FcRγ also supported expression of the CLR Mincle (*CLEC4E*) in T cells. Mincle is predominantly expressed by myeloid cells and recognizes trehalose-6,6′-dimycolate (TDM) from *Mycobacterium tuberculosis*, as well as fungal glycolipids^41–44^. When ectopically expressed alone in T cells, Mincle was not detected at the cell surface; however, co-expression with FcRγ resulted in robust surface expression in CD8⁺ and CD4⁺ T cells (Fig. 4k; Fig. S9e). Mincle-FcRγ co-expression enabled T cell activation in response to the Mincle agonist trehalose-6,6′-dibehenate (TDB), which was not observed with Mincle alone (Fig. 4l; Fig. S9f). Together, these findings extend FcRγ support to functional CLR expression, enabling conventional T cells to respond to microbial molecular patterns.

## Discussion

This study establishes FcRγ reconstitution as a generalizable approach for importing heterologous innate receptor functions into conventional T cells. Across natural cytotoxicity, Fc receptor, and C-type lectin families, FcRγ enabled or enhanced receptor surface expression and ligand-dependent T cell responses against novel cellular stimuli. Despite abundant endogenous CD3ζ, FcRγ played a non-redundant role in supporting the expression or function of the innate receptors tested.

Importantly, in T cells, FcRγ-associated receptors could engage conventional T cell signaling machinery despite differences in ITAM-based signaling components between innate immune cells and T cells. Using NKp30-FcRγ as an exemplar, we demonstrated ligand-dependent, TCR-like activation of canonical NFAT, NF-κB, and AP-1 transcriptional outputs in T cells, together with corresponding cytokine responses and cytotoxicity. Ligand-dependent functional responses observed across other FcRγ-associated receptor modules suggest that this compatibility extends across innate receptor families, reflecting the underlying homology between key components of innate and T cell ITAM-based signaling pathways^29,45,46^.

Modularity of FcRγ reconstitution offers a means to readily introduce different recognition modes into engineered conventional T cells. In our studies, these included expanded tumor recognition by NCR-T cells, enhanced ADCC and IgE-dependent activation by FcR-T cells, and microbial pattern recognition by CLR-T cells. These modules could plausibly complement existing TCR- and CAR-based therapies to mitigate tumor antigen escape or extend T cell applications to antimicrobial immunity.

Innate receptors normally operate within cellular contexts that integrate activating and inhibitory signals, and their transfer into conventional T cells may therefore alter receptor responsiveness, offering both therapeutic opportunities and potential safety challenges. The morbidity and tissue infiltration observed in mice treated with NKp46-FcRγ T cells highlight a need to assess receptor-specific tissue reactivity in the new T cell context and to define relevant ligands and ligand landscapes across human tissues as a prerequisite for clinical translation.

Together, our findings establish FcRγ reconstitution as a generalizable strategy for engineering conventional T cells with native innate receptors, thereby expanding their modes of recognition. Importantly, our phenotypic and transcriptional analyses demonstrate that FcRγ-enabled innate recognition does not require or induce an innate-like T cell state. This ability to combine diverse innate recognition modes with the effector functions and persistence of conventional T cells highlights the potential of FcRγ reconstitution for building upon this widely adopted engineered T cell therapy platform.

## Methods

### Cell lines and cell culture

Human cell lines were purchased from the American Type Culture Collection (ATCC) unless specified. The ovarian cancer cell line OVCAR-8 was generously provided by the Division of Cancer Treatment and Diagnosis (DCTD) Tumor Repository at the National Institutes of Health (NIH). All cells were maintained at 37°C and 5% CO₂ in a humidified incubator. Detailed cell line information and culture media are listed in Supplementary Data 1.

### Human peripheral blood mononuclear cells (PBMCs)

Human PBMCs from de-identified healthy adult blood donors were isolated by Ficoll-Paque (Cytiva) density gradient separation and obtained from the UCLA CFAR Virology Core. PBMCs were cryopreserved in aliquots of 1×10^7^ cells in CryoStor CS10 (Millipore Sigma) and thawed for each experiment.

### Primary human leukemia samples

Primary leukemia specimens were obtained under a University of California, Los Angeles (UCLA) Institutional Review Board (IRB)-approved protocol (IRB#22-000558). Bone marrow aspirates and peripheral blood were collected after informed consent from patients with AML or blast-phase MPN. Patient characteristics are summarized in Supplementary Table 1. Mononuclear cells were isolated by Ficoll-Paque density-gradient centrifugation, cryopreserved in CryoStor CS10 and thawed for each experiment.

### Plasmids

Lentiviral vector sequences used in this study are summarized in Supplementary Data 1. The full-length coding sequence of human B7-H6 (*NCR3LG1*, GenBank NM_001202439.3) was synthesized (Integrated DNA Technologies; IDT) and cloned into the second-generation lentiviral vector pCCL-c-MNDU3 (gift from Dr. Donald Kohn, UCLA) using Gibson Assembly (NEB). The ecto-calreticulin (ectoCRT) fusion protein was based on a published murine sequence (Addgene sequence-446903)^33^ and constructed by adding the GPI signal sequence from pRetroX-mCRT-GPI-IRES-DsRed in frame to the 3’ end of the full-length coding sequence of human calreticulin (*CALR*, GenBank NM_004343.4). The sequence was codon-optimized (GenScript), synthesized (IDT) and cloned into pCCL-c-MNDU3 by Gibson Assembly. The full-length coding sequences of human FcRγ (*FCER1G*, GenBank NM_004106.2), NKp30 (*NCR3*, GenBank NM_147130.3), NKp46 (*NCR1*, GenBank NM_004829), FcεRIα (*FCER1A*, GenBank NM_001387280.1), Dectin-2 (*CLEC6A*, GenBank NM_001007033.2), and Mincle (*CLEC4E*, GenBank NM_014358.4) were synthesized (IDT or Twist Bioscience) and cloned into pCCL-c-MNDU3 upstream of T2A and truncated *NGFR* sequences. For the CD16a construct, the full-length coding sequence of human CD16a (*FCGR3A*, higher-affinity 158V variant)^47^ was codon-optimized (GenScript), synthesized (IDT), and cloned into pCCL-c-MNDU3 upstream of T2A and truncated *NGFR* sequences. For all FcRγ co-expressing vectors, the stop codon was removed and a 3’ P2A sequence followed by full-length *FCER1G* was added; the construct was synthesized and cloned into the pCCL-c-MNDU3 vector with T2A-linked truncated *NGFR* as above.

For transcription factor reporter constructs, the NFAT-ZsGreen-mPGK-PuroR plasmid was a gift from Dr. David Baltimore (Caltech). The response elements (RE) for AP-1 and NF-kB were taken from pSIRV-AP-1-mCherry (Addgene) and pSIRV-NF-kB-eCFP (Addgene), respectively^48^, and used to replace the NFAT RE in NFAT-ZsGreen-mPGK-PuroR to generate AP-1-ZsGreen-mPGK-PuroR and NF-kB-ZsGreen-mPGK-PuroR vectors.

### Lentivirus preparation

Packaging and concentration of lentiviral particles were performed as previously described^49^ with certain modifications. Briefly, CHEDAR cells (gift from Dr. Donald Kohn, UCLA)^50^ were co-transfected with lentiviral vector plasmid, pCMV-ΔR8.9, and pCAGGS-VSVG using TransIT 293T (Mirus Bio) for 17 hours, followed by treatment with 20 mM sodium butyrate for 8 hours and generation of cell supernatants in serum-free Pro293a media (Lonza) supplemented with GlutaMax and 20 mM HEPES for 48 hours. Supernatants were concentrated by ultrafiltration using Amicon Ultra-15 100K filters (Millipore Sigma) at 4000 *g* for 40 minutes at 4°C and stored as aliquots at −80°C.

### T cell activation and transduction

Human T cells were isolated from cryopreserved peripheral blood mononuclear cells (PBMCs) by immunomagnetic depletion using the Pan-T Cell Isolation Kit (Miltenyi Biotec) and MACS LS Columns according to the manufacturer’s protocol, followed by activation with Dynabeads Human T-Activator CD3/CD28 (Thermo Fisher Scientific) at a 1:1 bead-to-cell ratio for 24 hours in AIM-V medium (Gibco) with 5% non-heat-inactivated human AB serum (Gemini Bio) (“A5” medium) supplemented with 100 IU/ml IL-2 (human IL-2 IS, Miltenyi Biotec). Dynabeads were then removed, and T cells were resuspended at 2×10^6^ cells/ml and plated on RetroNectin (Takara Bio) coated plates. Concentrated lentiviral supernatant was added together with 100 mg/ml poloxamer (Sigma-Aldrich) solution, for a final concentration of 1 mg/ml, and 100 IU/ml IL-2. At 5-6 days post-transduction, NGFR-expressing cells were purified using the MACSelect LNGFR System and LS Columns (Miltenyi Biotec). T cells were used for *in vitro* and *in vivo* experiments 6-11 days after transduction, unless specified.

### Primary NK cell isolation and expansion

Primary human NK cells were isolated from cryopreserved PBMCs by immunomagnetic depletion using the NK Cell Isolation Kit (Miltenyi Biotec) and MACS LS Columns. NK cells were activated and expanded by coculture with K562 cells transduced to express 4-1BBL (CD137L) and membrane-bound human IL-21 at an effector-to-target ratio of 1:2 for 2 weeks in RPMI 1640 media supplemented with 10% vol/vol FBS (“R10”), 1% L-glutamine, 1% penicillin/streptomycin, 50 uM 2-mercaptoethanol, 1 mM sodium pyruvate, 0.1 mM MEM nonessential amino acids, 10 mM HEPES, 20 ng/mL recombinant human IL-2 and 20 ng/mL recombinant human IL-15 (both from Thermo Fisher Scientific). NK cells were harvested on day 14 of expansion for *in vitro* and *in vivo* experiments.

### CRISPR-Cas9-mediated gene disruption

CRISPR-Cas9-mediated gene editing was performed by electroporation of sgRNA-Cas9 ribonucleoprotein (RNP) complexes using a 4D-Nucleofector X Unit (Lonza). For B7-H6 (*NCR3LG1*) knockout in K562 cells, 100 pmol sgRNA (Synthego) and 60 pmol Cas9 protein (QB3 MacroLab, University of California, Berkeley) were mixed and incubated at room temperature for 15 minutes for RNP complex assembly. 2×10^5^ K562 cells were resuspended in SF Nucleofector Solution (Lonza) containing the RNP complexes and electroporated using program FF-120. Five days after electroporation, B7-H6-negative cells were sorted by flow cytometry based on loss of cell-surface B7-H6 expression. For NKp30 (*NCR3*) knockout in primary T cells, FcRγ-transduced T cells were generated as described above. At 24 hours after lentiviral transduction, 1×10^6^ T cells were resuspended in P3 Primary Cell Nucleofector Solution (Lonza) containing RNP comple xes assembled from 100 pmol sgRNA and 60 pmol Cas9 protein and electroporated using program EH-115. Four days after electroporation, NGFR-positive, NKp30-negative T cells were sorted by flow cytometry, returned to culture and used for experiments the following day. For *TRAC* knockout in T cells, T cells were activated and transduced with lentivirus as described above. At 28-30 hours after transduction, 1×10^6^ T cells were resuspended in P3 Primary Cell Nucleofector Solution (Lonza) containing RNP complexes assembled from 200 pmol sgRNA and 100 pmol Cas9 protein and electroporated using program EH-115. T cells were subsequently expanded in A5/IL-2 as above and used for functional experiments 7-12 days after transduction. *TRAC* knockout efficiency was confirmed by loss of surface TCRαβ expression by flow cytometry. sgRNA target sequences are provided in Supplementary Data 1.

### Flow cytometry analysis

For surface marker staining, cells were incubated with Human TruStain FcX or TruStain FcX PLUS (BioLegend) to block Fc receptors. Cells were subsequently stained with fluorochrome-conjugated antibodies for 20 minutes at 4°C in the dark. All staining and washing steps were performed in PBS containing 0.5% bovine serum albumin (BSA) and 2 mM EDTA. For non-fixed samples, DAPI was added immediately before acquisition as a viability dye. For intracellular staining of NKp30, T cells were activated with anti-CD3/CD28 Dynabeads in the presence of 100 IU/ml IL-2 or 5 ng/ml IL-7 plus 20 ng/ml IL-15, as above, for 24 or 72 hours, followed by surface staining then fixation and permeabilization using Fixation and Permeabilization Wash buffers (BioLegend) according to the manufacturer’s protocol. Permeabilized cells were stained with an anti-NKp30 antibody (BioLegend) and analyzed by flow cytometry. Flow cytometry data were acquired using LSRFortessa (BD Biosciences), FACSCelesta (BD Biosciences) or Aurora (Cytek Biosciences) instruments and data analyzed using FlowJo software. Cell sorting was performed using FACSAria II or FACSAria H (BD Biosciences) instruments. A list of antibodies used for flow cytometry is provided in Supplementary Data 1.

### T cell cytokine withdrawal assays

For cytokine withdrawal experiments, 2 days post-transduction, T cells were washed twice with A5 medium and resuspended in A5 with or without 100 IU/ml IL-2 (human IL-2 IS, Miltenyi Biotec). Cells were expanded for an additional 8 days before flow cytometry analysis.

### T cell intracellular cytokine assays

2×10^5^ T cells were co-cultured with target cells at a 1:1 effector-to-target (E:T) ratio in 200 µl R10 without supplemented cytokines. For CD16a-dependent activation experiments, rituximab (1 µg/ml) or lintuzumab (1 µg/ml) was supplemented as indicated. Co-cultures were incubated for 6 hours in the presence of Protein Transport Inhibitor Cocktail (eBioscience). Positive control wells containing T cells only were stimulated with Cell Stimulation Cocktail (eBioscience), a cocktail of phorbol 12-myristate 13-acetate (PMA) and ionomycin. Following co-culture, cells were stained for surface markers and Zombie fixable viability dye (BioLegend) and fixed. Fixed cells were stored in Cyto-Last Buffer (BioLegend) at 4°C for up to 2 weeks, then washed, permeabilized, and stained for intracellular cytokines using the eBioscience Intracellular Fixation & Permeabilization Buffer Set (Thermo Fisher Scientific) according to the manufacturer’s two-step protocol for intracellular cytoplasmic proteins (Thermo Fisher Scientific BestProtocols).

### CD16 antibody stimulation

96-well plate wells were coated overnight with an anti-CD16 agonist antibody (clone CB16, Thermo Fisher Scientific) at 1:100 dilution in PBS and washed three times with PBS before T cell plating. T cells were resuspended in R10 supplemented with Protein Transport Inhibitor Cocktail (Thermo Fisher Scientific) and plated either on anti-CD16-coated wells or on untreated wells with soluble anti-CD16 (25 µg/ml) added to the media. After 6 hours, cells were harvested and processed for surface marker and intracellular cytokine staining as described above.

### T cell CD137 activation assays

For FcεRIα-mediated T cell activation assays, wells were coated with 10 μg/ml recombinant streptavidin in PBS (AcroBiosystems) for 3 hours at room temperature. Streptavidin-coated 96-well plate wells were then washed three times with PBS and coated with 10 μg/ml biotinylated human IgE-Fc Protein His Avitag (AcroBiosystems) in PBS for 1 hour at room temperature, then washed three times with PBS. 2×10^5^ T cells (FcRγ, FcεRIα, or FcεRIα-FcRγ T cells) were plated in streptavidin-only or streptavidin+IgE-coated wells in 200 μl A5 media without supplemented cytokines for 24 hours before flow cytometry analysis. For Dectin-2-mediated T cell activation, 2×10^5^ T cells (FcRγ, Dectin-2, or Dectin-2-FcRγ) were plated in a 96-well plate in 200 μl R10 with 0, 25, 50, 100, or 200 µg/ml furfurman (*Malassezia furfur* cell wall preparation; InvivoGen) and cultured for 18-24 hours before flow cytometry analysis. For Mincle-mediated T cell activation, 2×10^5^ T cells (FcRγ, Mincle, or Mincle-FcRγ) were plated in a 96-well plate in 200 μl R10 with 0, 1, 10, or 100 µg/ml trehalose-6,6-dibehenate (TDB) (Mincle agonist; InvivoGen) and cultured for 18-24 hours before flow cytometry analysis.

### T cell proliferation assays

1×10^5^ T cells were co-cultured at a 1:1 E:T ratio with 1×10^5^ target cells (Raji-mKate, Ramos-mKate, or NALM-6-mKate) in 200 µl R10 with 1 µg/ml rituximab (MedChemExpress) for 7 days in a 48-well plate. On day 4, media volume was doubled by addition of 200 µl R10. Cells were harvested at 7 days, stained with DAPI, and resuspended in 200 µl MACS buffer. A fixed volume was acquired on an Aurora (Cytek) spectral flow cytometer, and absolute viable cell counts were calculated based on the acquired volume using FlowJo software.

### Transcription factor (TF) reporter assays

T cells were isolated from total PBMCs and activated with Dynabeads as described above. At 24 hours post-activation, T cells were co-transduced with two lentiviral vectors encoding 1G4 TCR or NKp30-FcRγ, and ZsGreen under the control of NFAT, NF-κB, or AP-1 response elements. At 5 days post-transduction, MACS NGFR selection was performed to enrich for TCR- or NKp30-FcRγ-transduced T cells. NGFR-positive cells were then treated in culture with 1.25 µg/ml puromycin for 7 days to select for TF reporter-transduced cells. During the final two days of puromycin treatment, IL-2 was removed from the culture media to reduce background activation. T cells were then co-cultured with corresponding cognate K562 antigen-presenting cells (APCs): K562-A2/ESO SCT or K562-B7H6. T-cell-only wells treated with PMA/ionomycin were included as positive controls for each condition. ZsGreen reporter expression was analyzed at 24 hours by flow cytometry.

### Exhaustion-associated marker and cytokine recall assays

1G4 or NKp30-FcRγ T cells were activated, transduced, and enriched by MACS as described above. Transduced T cells were plated in A5 medium without supplemented cytokines and stimulated with corresponding cognate K562 APCs (either K562-A2/ESO SCT or K562-B7H6 cells) irradiated at 100 Gy. Irradiated APCs were added every 3 days at an E:T ratio of 1:1, 1:2, 1:4, or 1:8 for a total of 4 cycles. T cells were stained for exhaustion-associated markers before the first APC stimulation and 4 days after the final APC stimulation. T cells from the end of the same co-cultures were also stimulated with PMA/ionomycin in the presence of protein transport inhibitors and analyzed by intracellular cytokine staining as described above.

### IncuCyte cytotoxicity and serial antigen rechallenge assays

Fluorescent target cell lines were established by lentiviral transduction with mKate2xNLS (nuclear-localized mKate) based on a published sequence^51^. Tissue culture-treated, black-walled, flat-bottom 96-well plates (3904, Corning) were coated with Poly-L-Lysine (Millipore Sigma) for 1 hour at room temperature and allowed to dry at room temperature for 2 hours. Target cells (1-2×10^4^ cells, depending on the cell line) were plated in 100 µl R10 without supplemented cytokines either the afternoon before (for adherent cell lines) or one hour before (for suspension cells) addition of effector cells. T cells or NK cells were added at a 1:1 E:T ratio in 100 µl R10, unless specified. For CD16a experiments, rituximab (1 µg/ml) or lintuzumab (1 µg/ml) was supplemented accordingly. For serial antigen rechallenge experiments, 100 µl supernatant was carefully removed without disturbing the cells every 3 to 4 days, and fresh target cells were added in 100 µl R10 at the original target cell seeding number. All IncuCyte experiments were set up with triplicate wells for each condition. Target-cell abundance was quantified every 2 hours for 5-20 days using an IncuCyte SX5 Live-Cell Imaging System (Sartorius). Target-cell abundance was assessed by total red object counts.

### Primary acute myeloid leukemia (AML) killing assays

AML patient specimens were selected based on blast CD34+ positivity. CD34+ AML cells were isolated from these samples by MACS using the UltraPure CD34 MicroBead Kit and LS Columns (Miltenyi Biotec). CD34-enriched AML cells were then labeled with CellTrace Violet (Thermo Fisher Scientific) according to the manufacturer’s protocol and co-cultured at an E:T ratio of 1:1 with *TRAC*-knockout NKp30-FcRγ or NGFR control T cells prepared as described above. Flow cytometry staining with Apotracker Green (BioLegend) and DAPI was performed after 72 hours of coculture to quantify apoptosis. For analysis, leukemia cells were distinguished from effector T cells by CTV positivity. For primary AML IncuCyte experiments, CD34+ AML blasts were cultured in X-VIVO-15 (Lonza) media supplemented with SCF, FLT3L, TPO, and IL-3 (20 ng/ml each) and transduced, as described above, with mKate2xNLS and, after 4 days, used in IncuCyte assays with *TRAC*-knockout NKp30-FcRγ or NGFR control T cells in R10 media, as described above.

### Western blot analysis of tumor cell lines and primary AML cells

Cell lines or isolated CD34+ primary AML cells were lysed with RIPA buffer supplemented with protease and phosphatase inhibitors. Lysates were incubated on ice for 15 minutes and vortexed at 3-minute intervals. Cell debris was pelleted by centrifugation at 12,000 *g* for 12 minutes at 4°C, and supernatants were transferred to low-protein-binding tubes. Protein concentrations were determined by bicinchoninic acid (BCA) assay using the Pierce Dilution-Free Rapid Gold BCA Protein Assay Kit (Thermo Fisher Scientific). Protein lysates were diluted with LDS sample buffer and reducing agent and denatured at 70°C for 10 minutes. 15 µg protein per sample was resolved by SDS-PAGE using 4-12% Bis-Tris gels with MES SDS running buffer and subsequently transferred to polyvinylidene fluoride (PVDF) membranes using an iBlot 3 Western Blot Transfer Device (Thermo Fisher Scientific). Transferred membranes were blocked in 5% non-fat milk in TBST for 30 minutes at room temperature and washed 3 times in TBST, followed by incubation overnight at 4°C with agitation in primary antibodies diluted in 5% non-fat milk in TBST. Primary antibodies used were B7-H6 (E9B2L), β-actin (13E5), and GAPDH (14C10) rabbit monoclonal antibodies (all from Cell Signaling Technology). Membranes were subsequently washed 3 times in TBST and incubated with horseradish peroxidase (HRP)-conjugated anti-rabbit secondary antibody (Cell Signaling Technology) for 1 hour at room temperature with agitation. All antibodies were used at 1:1000 dilution. Bound antibody was visualized using SuperSignal West Pico PLUS chemiluminescent substrate (Thermo Fisher Scientific) and imaged using a ChemiDoc MP Imaging System (Bio-Rad). β-actin or GAPDH staining was used as a loading control.

### Single-cell RNA-sequencing

PBMCs from two healthy donors were activated and cells from each donor were either transduced with FcRγ-NGFR lentivirus or non-transduced. At 5 days post-transduction, NGFR-positive total T cells or non-transduced total T cells were isolated by FACS and resuspended in PBS/0.04% BSA. 10X Chromium 5’ GEX libraries were constructed using the 10X Genomics Chromium Controller Single Cell Sequencing System (10X Genomics) by the UCLA TCGB core facility. Libraries were sequenced on a NovaSeq X Plus 25B (Illumina) instrument. Single-cell RNA-seq data were processed and aggregated using Cell Ranger software (10x Genomics), and reads were aligned to a human reference genome assembly. Gene expression counts were generated using the Cell Ranger “Count” pipeline. Cell clustering, UMAP visualization, feature plots of selected genes, and pseudobulk differential gene expression analysis were performed using Loupe Browser (10x Genomics).

### Mice

Female NOD.Cg-*Prkdcscid Il2rgtm1Wjl*/SzJ (NSG) mice at 6-8 weeks of age were obtained from The Jackson Laboratory and housed at the California NanoSystems Institute vivarium at UCLA. All animal experiments were performed in accordance with guidelines established by the UCLA Division of Laboratory Animal Medicine (DLAM) under protocols approved by the UCLA Animal Research Committee (ARC).

### *In vivo* tumor xenograft models

Mouse experiments were performed under a protocol approved by the UCLA Animal Research Committee (ARC). All mice were purchased from The Jackson Laboratory. HL-60 AML xenograft mice were established by injecting female NSG mice, 6-8 weeks of age, intravenously via tail vein with 5×10^5^ HL-60 cells stably transduced with firefly luciferase and IRES-linked GFP (HL60-luc-GFP). For T cell efficacy studies, 3 days after tumor cell injection, tumor burden was assessed by intraperitoneal (IP) luciferin injection and bioluminescence imaging (BLI) performed at the Crump Preclinical Imaging Technology Center at UCLA. Mice were then allocated to treatment groups based on BLI to achieve comparable mean tumor burden across groups. 10 mice per group each received PBS or 1×10^7^ NGFR control T cells and 12 mice received 1×10^7^ NKp30-FcRγ T cells by tail vein injection. The day of adoptive immune cell transfer was defined as day(D) 0. BLI was performed starting on D+4 and twice weekly thereafter. Mice were sacrificed based on morbidity criteria per institutional guidelines. At experimental endpoints, blood was obtained by retro-orbital collection into heparinized capillary tubes for flow analysis. Spleen and bone marrow were collected for flow analysis, and the indicated tissues were prepared for histological and immunohistochemical analyses as described below. T cell/NK cell comparison experiments were performed similarly, with modifications for stringency: 4 days after tumor cell injection, 5×10^6^ T cells or 5×10^6^ NK cells were injected per mouse on D0, and BLI commenced on D+3. For the NALM-6 tumor model, NSG mice were intravenously injected via tail vein with 1×10^5^ NALM-6-luc-GFP cells. Four days after tumor cell injection, tumor burden was assessed by BLI and mice were allocated to treatment groups to achieve comparable mean tumor burden across groups. On D0, mice received 5×10^6^ non-transduced (NTD) or an equal number of NKp46-FcRγ T cells via tail vein (*n*=10 mice per group). BLI and endpoint analyses were performed as above.

### Histology and immunohistochemistry (IHC)

Mice were sacrificed based on morbidity criteria per institutional guidelines, and relevant organs were embedded in Tissue-Tek O.C.T. Compound (Andwin Scientific) and frozen on dry ice. Immunohistochemical detection of CD3 was performed by the UCLA Translational Pathology Core Laboratory (TPCL). Briefly, frozen tissues were sectioned at a thickness of 4 µm and stained using an automated BOND immunostaining system. Sections underwent heat-induced epitope retrieval (HIER) in ER2 retrieval solution for 20 min at 100°C, followed by a wash step, and endogenous peroxidase activity was blocked for 5 min. Sections were then incubated with DAKO polyclonal rabbit anti-human CD3 antibody (catalog no. A0452; Agilent/Dako) at a 1:100 dilution for 15 min, followed by a polymer-based secondary detection reagent for 8 min. Immunoreactivity was visualized using Mixed DAB Refine for 10 min, followed by hematoxylin counterstaining for 5 min. Washes were performed between staining steps according to the automated BOND protocol. The A0452 antibody is an affinity-isolated polyclonal rabbit anti-human CD3 reagent that is also specified for use on frozen sections. Images were acquired using an Olympus BX51 microscope

## Data Availability

Single-cell RNA-sequencing datasets generated during the current study will be made available in a public repository upon final manuscript publication.

## Acknowledgements

We thank the members of the Seet Lab for their valuable support. We thank Felicia Codrea, Jessica Scholes, and Jeff Calimlim of the Broad Stem Cell Research Center Flow Cytometry Core for assistance with cell sorting; Melody Li and Xinmin Li of the UCLA TCGB Core for assistance with 10X sequencing; Profs. Gay Crooks, Steven Dubinett, Donald Kohn, Caroline Kuo, Sanaz Memarzedeh, Cristina Puig-Saus, Ramin Salehi-Rad, Anthony Wang, Owen Witte (UCLA), Joshua Sasine (Cedars-Sinai), Dr. Michael Bethune and the late Prof. David Baltimore (Caltech) for generously providing cell lines and plasmids; and the UCLA Broad Stem Cell Research Center for essential equipment support.

## Author Contributions

C.H. and S.L. designed, performed, and supervised experiments, analyzed data, and contributed to manuscript writing and preparation. Y.N. designed and performed key experiments and analyzed data. J.P. performed experiments and oversaw *in vivo* studies. J.C., K.C., and W.H. performed and assisted with experiments. K.D., O.Z., J.H., J.C.K.W., N.S., and V.C. assisted with experiments.

S.M. contributed valuable cell lines. C.O. consented patients and contributed patient specimens.

S.P. developed the patient specimen collection protocol, consented patients, contributed patient specimens, and assisted with experiments. A.M-H. and J.T.K. collaborated on experimental design and contributed key reagents. C.S.S. conceptualized, designed, and supervised experiments, analyzed data, and developed the manuscript.

---

**Fig S1:**
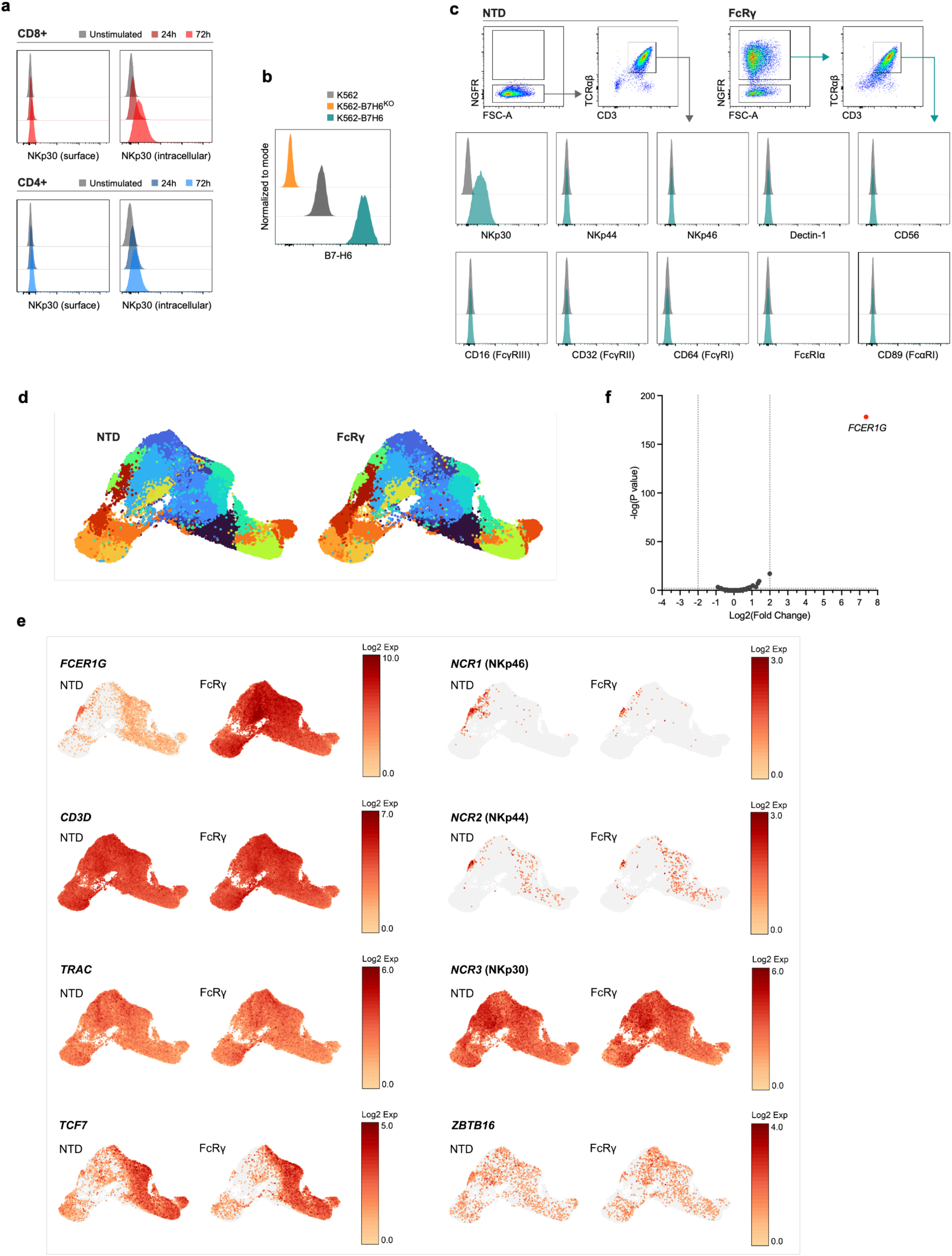
FcRγ enables functional endogenous NKp30 surface expression in activated conventional T cells. **a)** Flow cytometry expression of surface or intracellular NKp30 in CD8+ and CD4+ T cells activated with anti-CD3/28 beads and IL-7/IL-15 for the indicated times. Relative frequencies shown are normalized to mode. **b)** Flow cytometry expression of B7-H6 on parental K562, K562-B7H6^KO^, or K562-B7H6 (B7-H6 overexpressing) cell lines. Relative frequencies shown are normalized to mode. **c)** Surface expression of NK- and FcRγ-associated surface proteins on non-transduced (NTD) or FcRγ (with NGFR transduction marker) transduced total T cells (linked to NGFR marker). Histograms show relative frequencies normalized to mode. **d)** Single-cell RNA-seq gene expression clusters in NTD or FcRγ-transduced total T cells. Aggregated data from two independent PBMC donors is shown. **e)** Expression of canonical T cell or NK/innate-like T cell genes in NTD or FcRγ-transduced total T cells. Aggregated data from two independent PBMC donors is shown. **f)** Volcano plot of differentially expressed genes between NTD or FcRγ-transduced total T cells. Aggregated data from two independent PBMC donors is shown.

**Fig S2:**
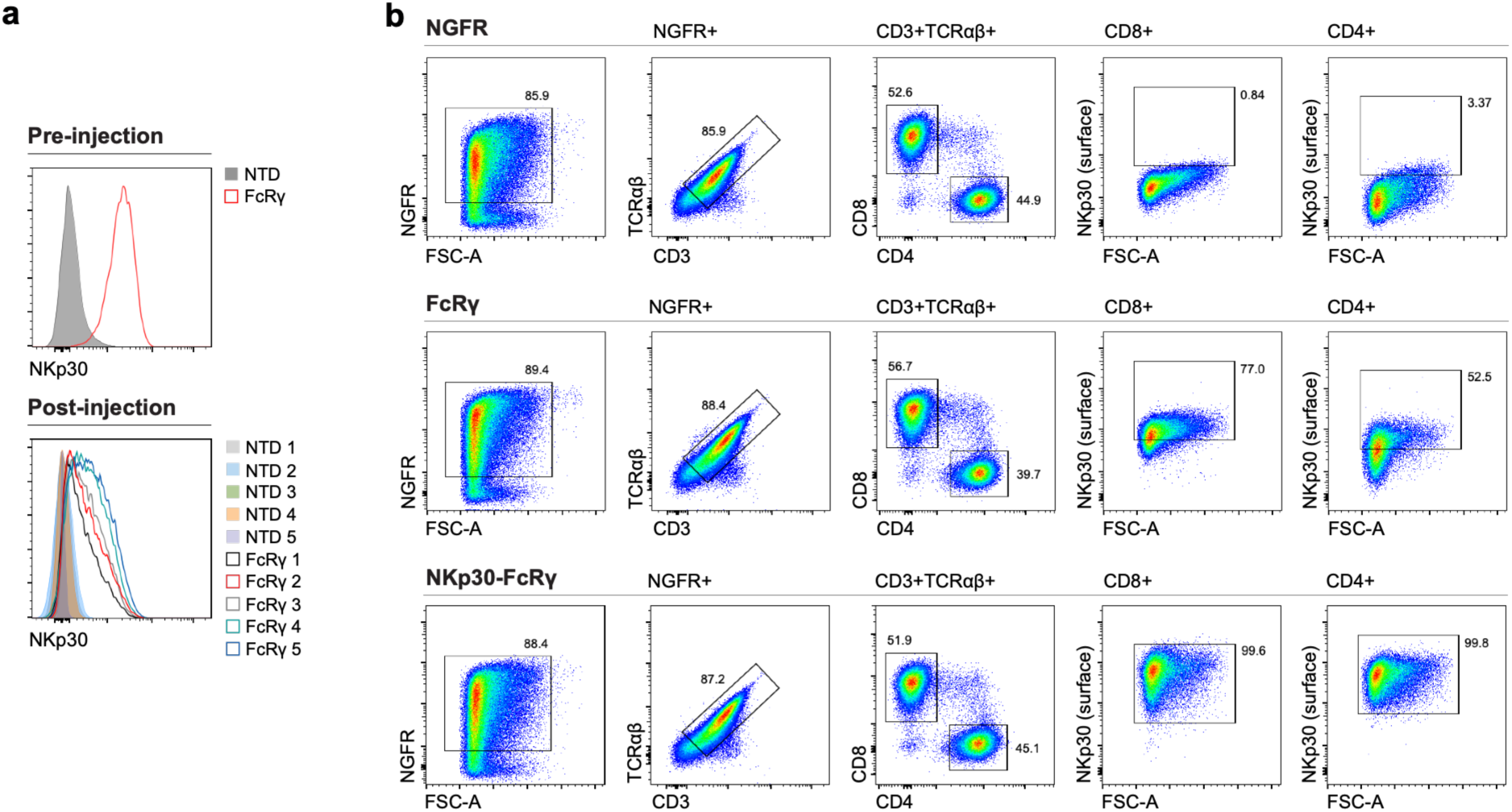
NKp30-FcRγ engineering creates stable, ligand-gated NCR-T cells. **a)** Surface NKp30 expression on non-transduced (NTD) or FcRγ-T cells before transplantation into NSG mice or 34-64 days post-transplantation (at the experimental endpoint for each mouse). Peripheral blood flow cytometry data is shown (*n*=5 mice per group). Relative frequencies shown are normalized to mode. **b)** NKp30 expression and phenotype of CD8+ and CD4+ T cells transduced with NGFR-only (empty vector control), FcRγ alone, or NKp30 with FcRγ (NKp30-FcRγ). Sequential gating is shown.

**Fig S3:**
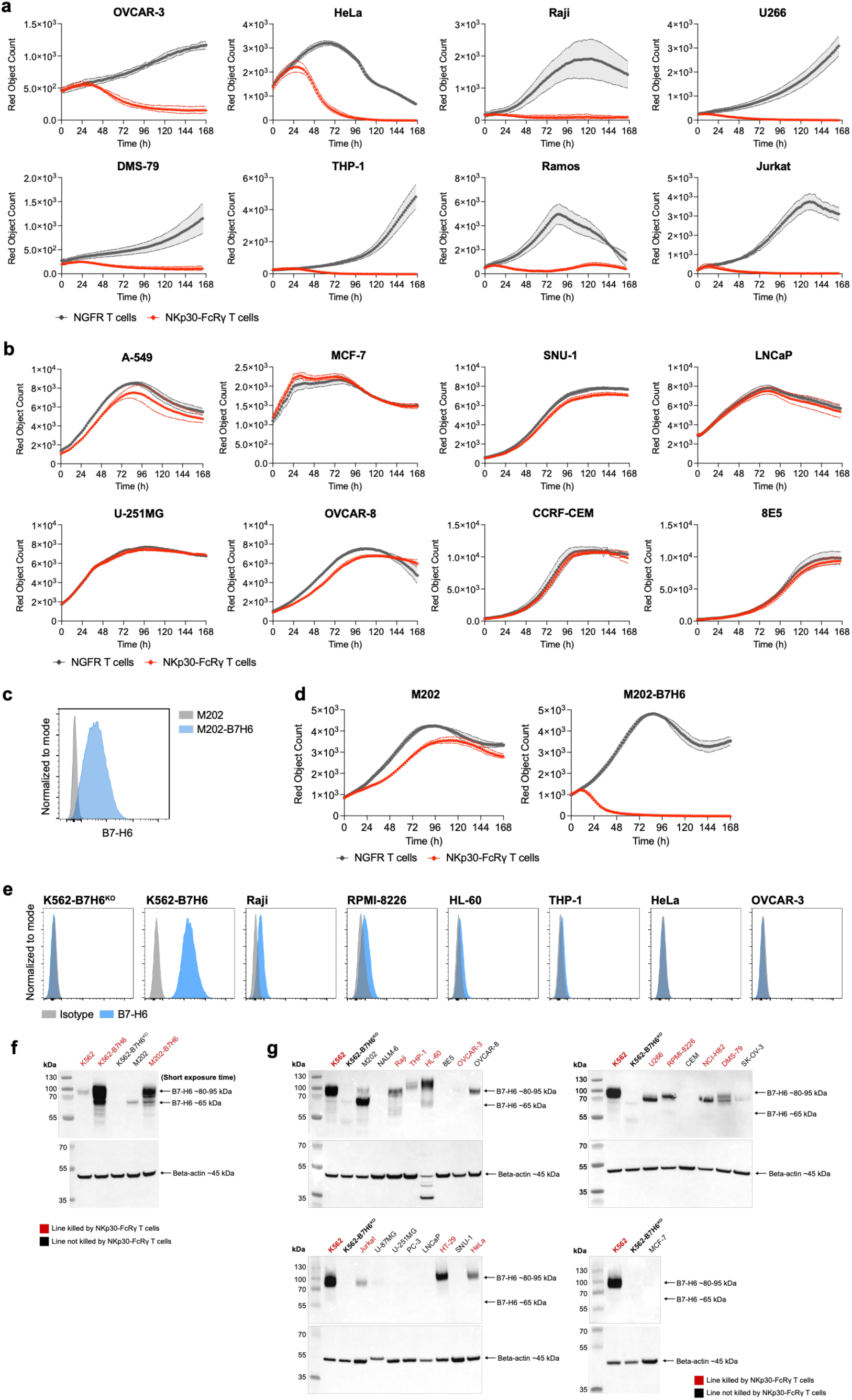
Stable NKp30–FcRγ engineering creates NCR-T cells with anti-tumor activity. **a)** Continued from Fig. 2d. IncuCyte counts of mKate-transduced tumor cell lines (described in Supplementary Data 1) co-cultured with *TRAC*^KO^ NGFR-only or *TRAC*^KO^ NKp30-FcRγ total NCR-T cells at a 1:1 E:T without exogenous cytokines. Representative cell lines killed by NKp30-FcRγ NCR-T cells are shown. Data are mean ± SEM of triplicate wells. **b)** Continued from Fig. 2e. IncuCyte assays, as described in panel a, for representative cell lines not killed by NKp30-FcRγ NCR-T cells. Data are mean ± SEM of triplicate wells. **c)** Expression of B7-H6 on parental M202 melanoma or M202-B7H6 (B7-H6 overexpressing) cell lines. Relative frequencies shown are normalized to mode. **d)** IncuCyte counts of mKate-transduced M202 or M202-B7H6 cells co-cultured with total T cells transduced with NGFR-only or NKp30-FcRγ NCR-T cells at a 1:1 E:T without exogenous cytokines. Data are mean ± SEM of triplicate wells. **e)** Flow cytometry surface expression of B7-H6 on selected cell lines (described in Supplemental Data 1). Relative frequencies shown are normalized to mode. **f)** Western blots of whole cell lysates of K562 and M202 cell lines and their B7H6^KO^ or overexpression derivatives. Arrows indicate reported B7-H6 molecular weights^24^. Exposure was decreased relative to blots shown in panel g to visualize high B7-H6 levels in the overexpressing cell lines. Beta-actin staining is shown as a loading control. Protein ladder is shown in the left lane of each blot. **g)** Western blots of whole cell lysates of a panel of tumor cell lines. K562 and K562-B7H6^KO^ cells are included on each blot as staining controls. Cell line names in red indicate lines killed by NKp30-FcRγ NCR-T cells in IncuCyte assays (Fig. 2d and Fig. S3a). Arrows indicate known B7-H6 molecular weights^24^. Beta-actin staining is shown as a loading control. Protein ladder is shown in the left lane of each blot.

**Fig S4:**
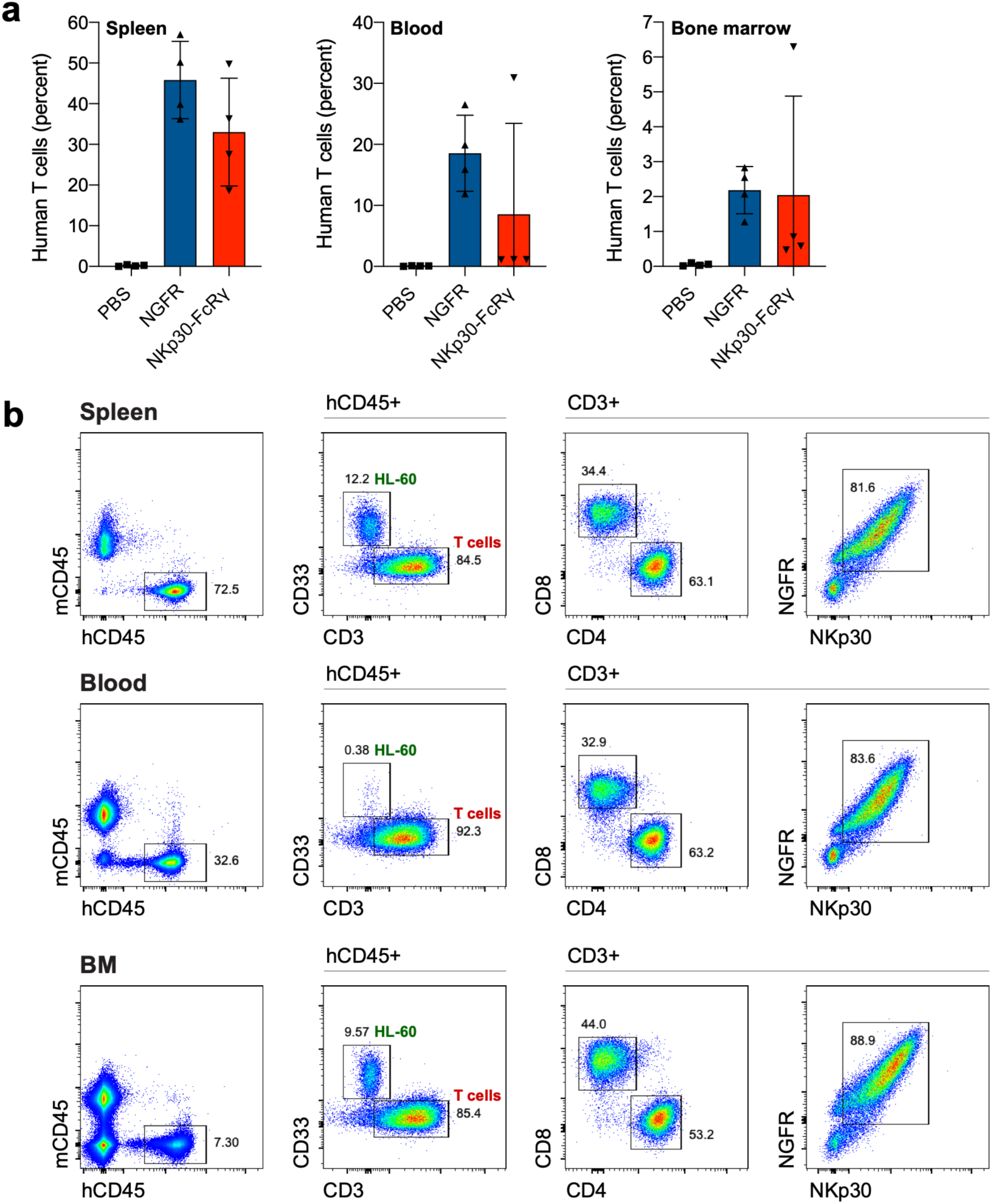
Stable NKp30-FcRγ engineering creates NCR-T cells with *in vivo* anti-leukemic activity. **a)** Necropsy analysis of mice described in Fig. 2f,g,h at the time of tumor progression. Frequency of engineered human T cells (CD3+NGFR+) as a percent of total cells in the spleen, blood, or bone marrow of recipient mice. Error bars represent the mean ± SD of four mice analyzed per experimental cohort. **b)** Representative flow cytometry phenotype of T cells from the tissues of mice receiving NKp30-FcRγ NCR-T cells at the time of tumor progression and death. Sequential gates are indicated.

**Fig S5:**
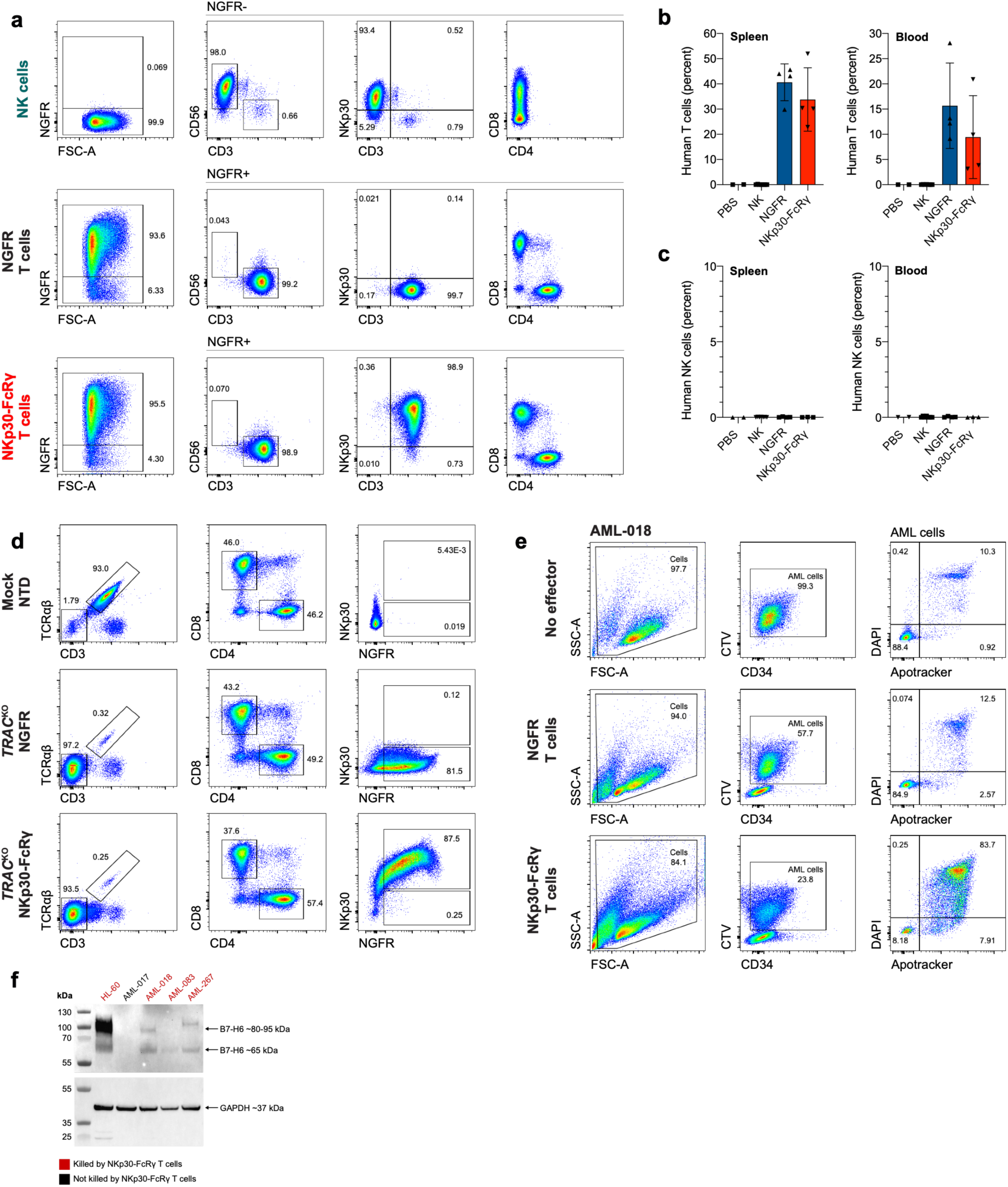
NKp30-FcRγ NCR-T cells show sustained activity compared with primary NK cells and target primary AML. **a)** Flow cytometry phenotypes and surface NKp30 expression of primary NK cells expanded with K562-mbIL21 cells and IL-15/IL-2, versus same-donor total T cells transduced with NGFR (empty vector control) or NKp30-FcRγ. **b)** Necropsy analysis of mice described in Fig. 3c at the time of tumor progression. Frequency of engineered human T cells (CD3+NGFR+) as a percent of total cells in the spleen or blood of recipient. Error bars represent the mean ± SD of 2-7 mice analyzed per experimental cohort. **c)** Necropsy analysis of mice described in Fig. 3c at the time of tumor progression. Frequency of engineered human NK cells (CD3-CD56+) as a percent of total cells in the spleen or blood of recipient. Error bars represent the mean ± SD of 2-7 mice analyzed per experimental cohort. **d)** Representative experiment showing flow cytometry phenotype of total T cells after *TRAC* deletion (TRAC^KO^) and non-transduced (NTD) or transduced with NGFR (empty vector control) or NKp30-FcRγ. **e)** Representative experiment showing gating strategy for primary AML apoptosis assay shown in Fig. 3e. CD34-selected AML blast cells were labeled with CellTrace Violet (CTV) prior to T cell co-culture. Sequential gating is shown. Apotracker labels apoptotic cells and DAPI labels dead cells. **f)** Western blot for B7-H6 expression in whole cell lysates from HL-60 cells (positive staining control) or primary AML blasts shown in Fig. 3e. Cell names in red indicate samples that were killed by NKp30-FcRγ NCR-T cells in Fig. 3e. Arrows indicate reported B7-H6 molecular weights^24^. GAPDH staining is shown as a loading control. Protein ladder is shown in the left lane of each blot.

**Fig S6:**
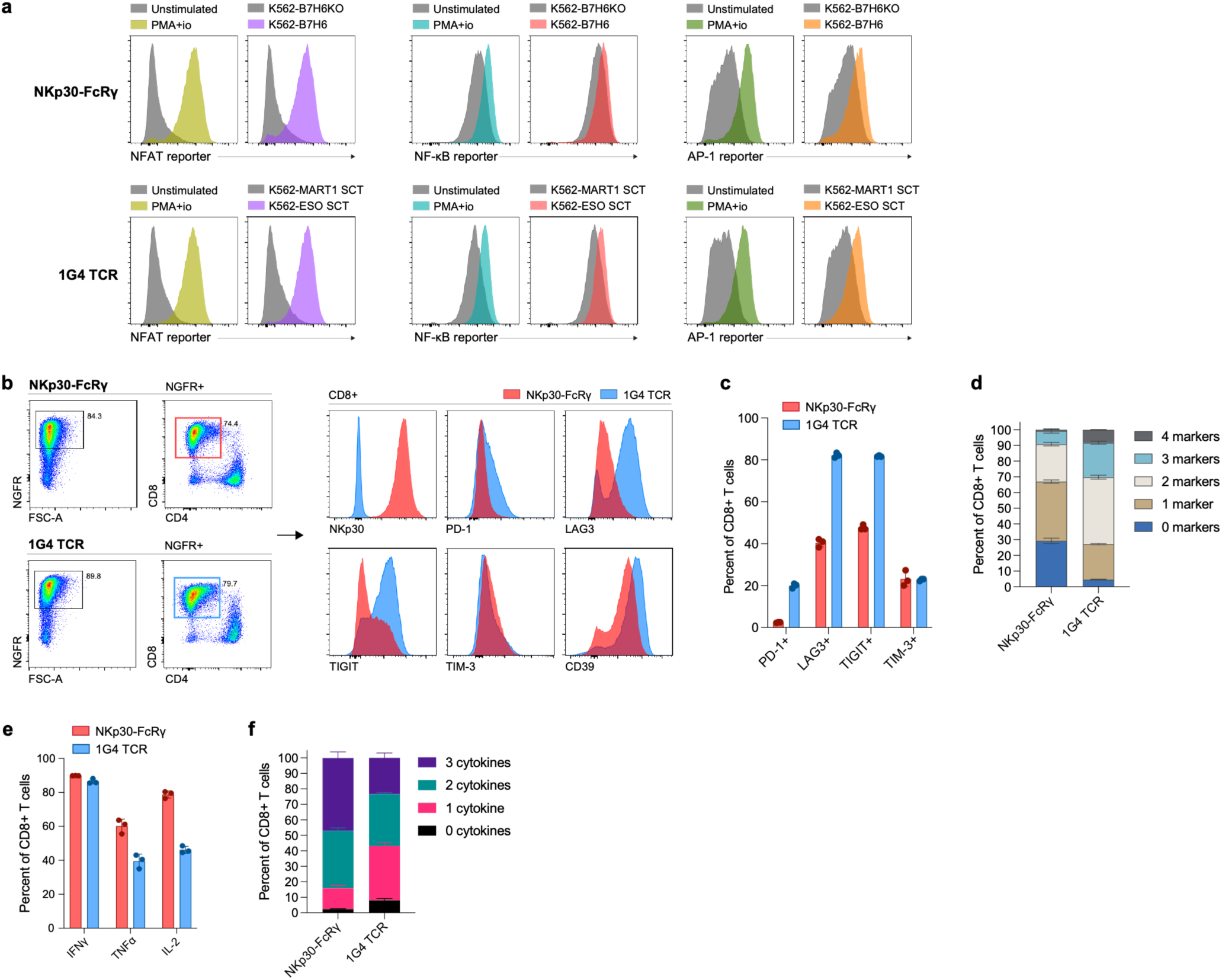
NKp30-FcRγ engagement couples innate recognition to TCR signaling outputs. **a)** Expression of ZsGreen driven by NFAT, NF-κB, or AP-1 response elements in primary T cells transduced with NKp30-FcRγ or 1G4 TCR (recognizing HLA-A*02:01/NY-ESO-1_157-165_). Reporter-transduced T cells were co-cultured for 24 hours with antigen-negative or antigen-positive target cells: K562-B7H6^KO^ or K562-B7H6, respectively, for NKp30-FcRγ NCR-T cells; and K562-MART1 SCT or K562-ESO SCT, respectively, for 1G4 TCR-T cells. T cell-only conditions with or without PMA/ionomycin served as positive and negative controls, respectively. Relative frequencies shown are normalized to mode. **b)** Expression of exhaustion-associated markers on CD8+ NKp30-FcRγ NCR-T or 1G4 TCR-T cells after 4 serial 3-day stimulations with cognate K562 target cells (K562-B7H6 or K562-ESO SCT, respectively). Relative frequencies shown are normalized to mode. **c)** Frequency of CD8+ T cells, as described in panel b, expressing each exhaustion-associated marker. Error bars represent mean ± SD of technical triplicate wells. **d)** Frequency of CD8+ T cells, as described in panel b, expressing Boolean combinations of PD-1, TIGIT, TIM-3, or LAG3. Error bars represent mean ± SD of technical triplicate wells. **e)** Frequency of CD8+ T cells, as described in panel b, producing IFNγ, TNFα, or IL-2, assessed by intracellular flow cytometry, in response to treatment with PMA/ionomycin for 6 h. Error bars represent mean ± SD of technical triplicate wells. **f)** Frequency of CD8+ T cells, as described in panel b, producing Boolean combinations of the cytokines IFNγ, TNFα, or IL-2 by intracellular flow cytometry in response to treatment with PMA/ionomycin for 6 h. Error bars represent mean ± SD of technical triplicate wells.

**Fig S7:**
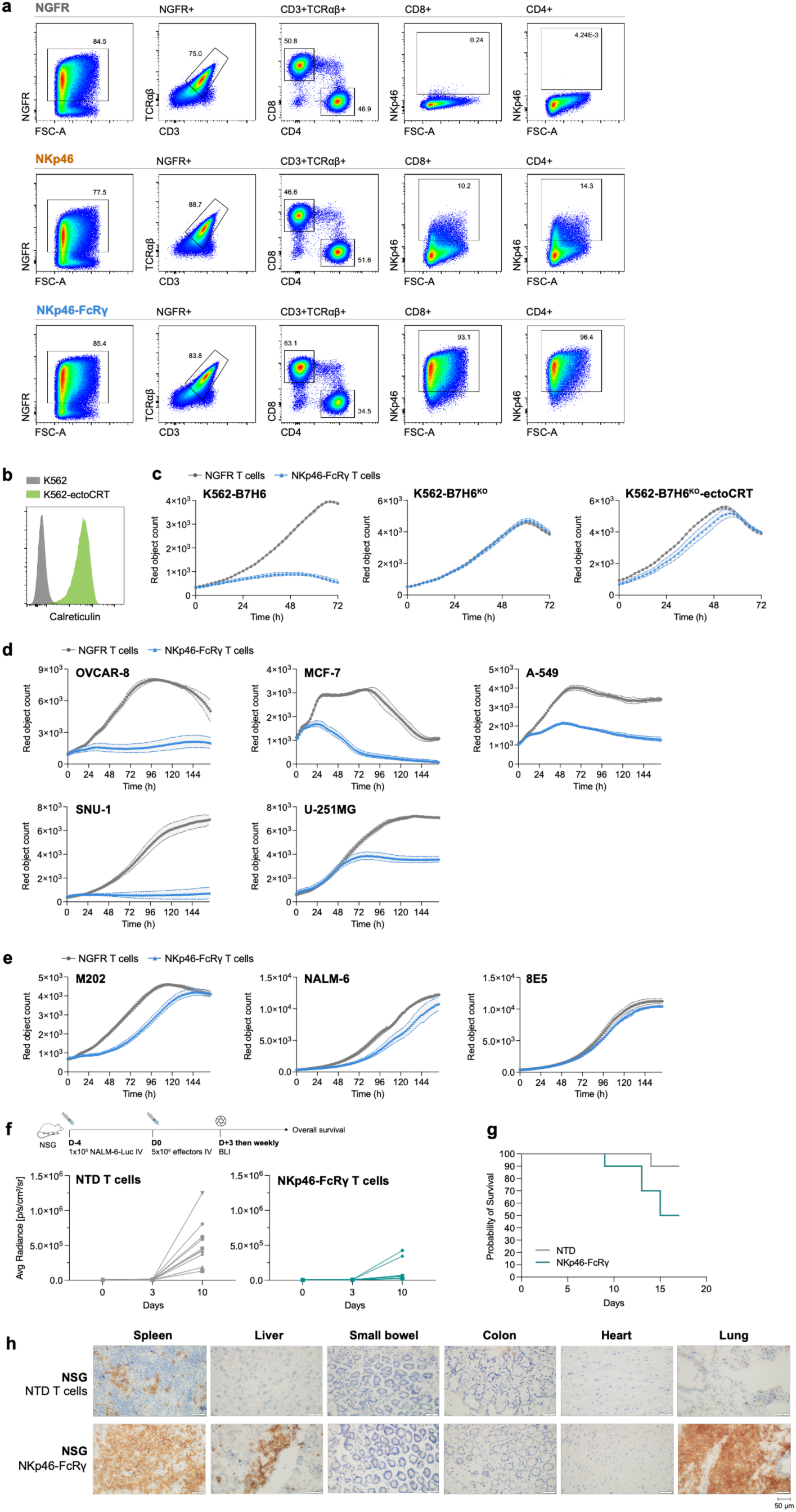
NKp46-FcRγ extends tumor recognition and reveals context-dependent reactivity. **a)** Flow cytometry phenotypes and surface NKp46 expression of T cells transduced with NGFR (empty vector control), NKp46, or NKp46-FcRγ. Sequential gating is shown. **b)** Stable surface expression of a GPI-calreticulin fusion protein (ectoCRT) on transduced K562 cells. **c)** IncuCyte counts of mKate-transduced K562-B7H6, K562-B7H6^KO^, or K562-B7H6^KO^-ectoCRT cells co-cultured with total NGFR T cells or NKp46-FcRγ NCR-T cells at a 1:1 E:T without exogenous cytokines. Data are mean ± SEM of triplicate wells. **d)** Continued from Fig. 4c. IncuCyte counts of mKate-transduced tumor cell lines co-cultured with *TRAC*^KO^ NGFR-only or *TRAC*^KO^ NKp46-FcRγ total NCR-T cells at a 1:1 E:T without exogenous cytokines. Data are mean ± SEM of triplicate wells. Representative cell lines killed by NKp46-FcRγ NCR-T cells are shown. **e)** Continued from Fig. 4C. IncuCyte assays as shown in panel d; cell lines partially killed by NKp46-FcRγ NCR-T cells are shown. **f)** Schematic of NALM-6 xenograft/adoptive cell therapy NSG mouse model and tumor response plots of luciferase bioluminescence imaging of NALM-6-luciferase engrafted mice which received total T cells which were either non-transduced (NTD) or transduced with NKp46-FcRγ (*n*=10 per group). **g)** Kaplan-Meier survival probability curves of mice cohorts shown in panel f. **h)** Immunohistochemistry staining for human CD3 in necropsy specimens from mice shown in panel f that received NTD or NKp46-FcRγ NCR-T cells. Tissue sections from one representative mouse per group are shown.

**Fig S8:**
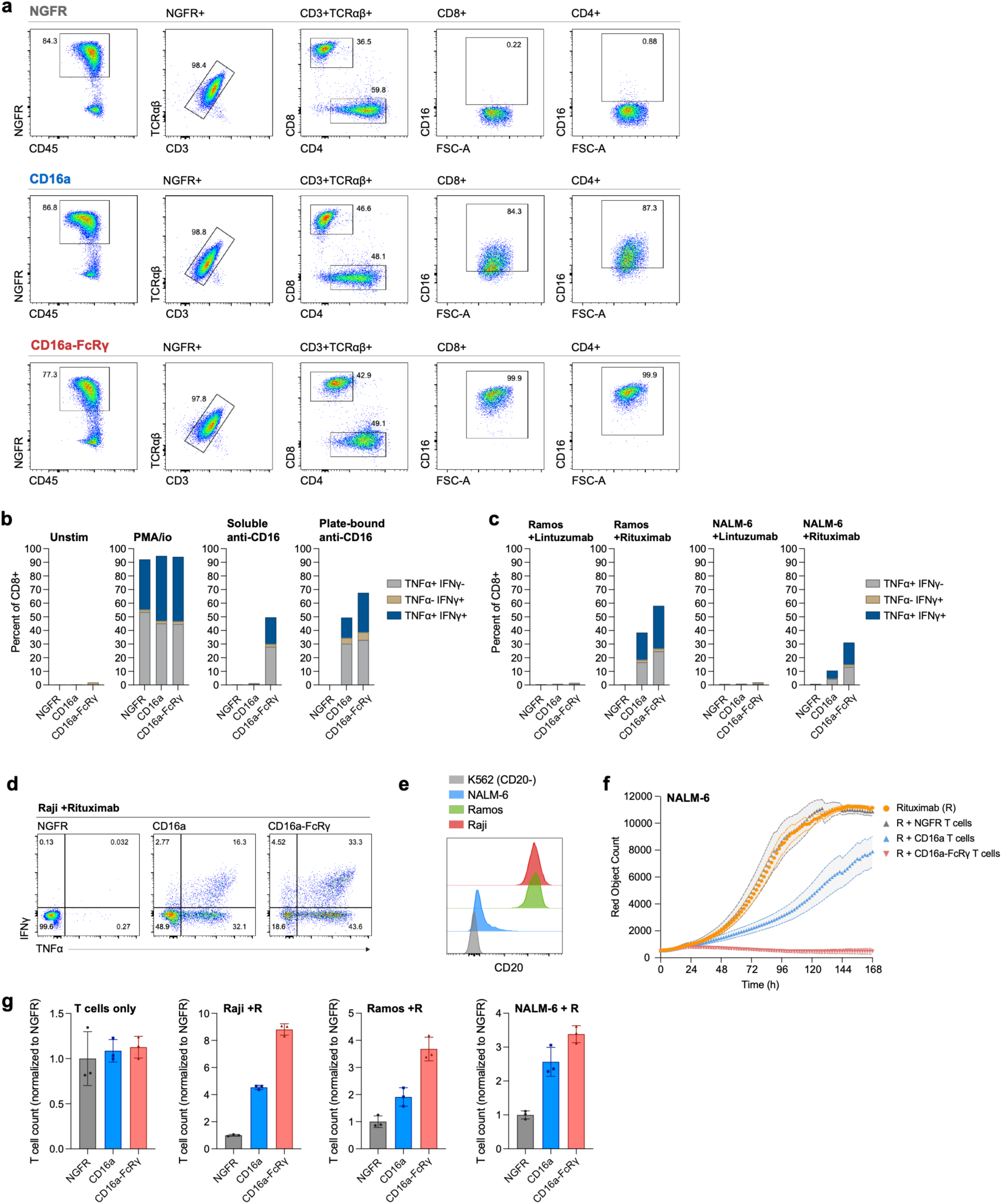
FcRγ improves antibody-dependent cytotoxicity in T cells through CD16a. **a)** Flow cytometry phenotypes and surface CD16a expression of T cells transduced with NGFR (empty vector control), CD16a or CD16a-FcRγ. **b)** Continued from Fig 4e. Intracellular IFNγ and TNFα expression in CD8+ T cells transduced with NGFR, CD16a, or CD16a-FcRγ unstimulated, treated with PMA and ionomycin (positive control), or either soluble or plate-bound anti-CD16 for 6 hours. **c)** Continued from Fig 4e. Intracellular IFNγ and TNFα expression in CD8+ T cells transduced with NGFR, CD16a, or CD16a-FcRγ co-cultured with CD20^high^ Ramos cells or CD20^low^ NALM-6 cells for 6h in the presence of Lintuzumab (anti-CD33) or Rituximab (anti-CD20) antibodies. **d)** Representative flow cytometry plots of intracellular IFNγ and TNFα expression in CD8+ T cells transduced with NGFR (control), CD16a or CD16a-FcRγ in response to 6h co-culture with CD20^high^ Raji cells in the presence of 1 µg/ml of Rituximab. **e)** Endogenous expression levels of CD20 on K562, NALM-6, Ramos, and Raji cell lines. **f)** IncuCyte counts of mKate-transduced NALM-6 cells co-cultured with NGFR, CD16a, or CD16a-FcRγ T cells at E:T 1:2 without exogenous cytokines in the presence of 1 µg/ml Rituximab (R). Data are mean ± SD of technical triplicate samples. **g)** *In vitro* expansion of NGFR, CD16a, or CD16a-FcRγ T cells co-cultured with Raji, Ramos, or NALM-6 cells in the presence of Rituximab (R) for 7 days. Error bars represent mean ± SD of technical triplicate wells; individual wells are shown as points. Values were normalized to the NGFR (empty vector) control.

**Fig S9:**
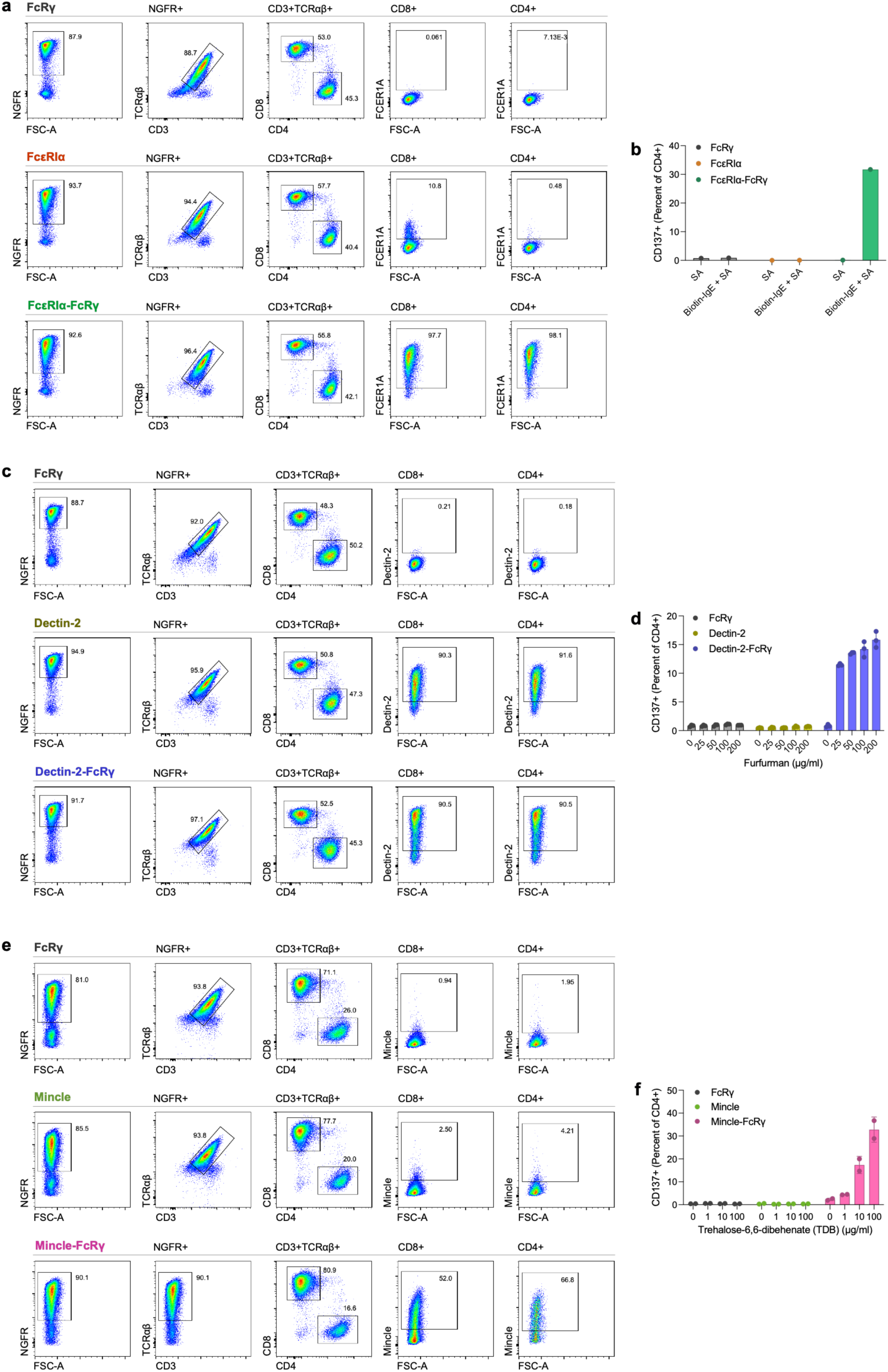
FcRγ supports FcR and CLR receptor modules in conventional T cells. **a)** Flow cytometry phenotypes and surface FcεRIα expression of T cells transduced with FcRγ alone, FcεRIα or FcεRIα-FcRγ (with NGFR marker). **b)** Continued from Fig 4h. Activation-induced CD137 expression on CD4+ T cells transduced with FcRγ alone, FcεRIα, or FcεRIα with FcRγ cultured for 24h with or without 10 μg/ml biotinylated IgE-Fc and streptavidin (SA). **c)** Flow cytometry phenotypes and surface Dectin-2 expression of T cells transduced with FcRγ alone, Dectin-2 or Dectin-2-FcRγ (with NGFR marker). **d)** Continued from Fig 4j. Activation-induced CD137 expression on CD4+ T cells transduced with FcRγ alone, Dectin-2, or Dectin-2 with FcRγ cultured for 18 h with furfurman at the indicated concentrations. **e)** Flow cytometry phenotypes and surface Mincle (CLEC4E) expression of T cells transduced with FcRγ alone, Mincle or Mincle-FcRγ (with NGFR marker). **f)** Continued from Fig 4l. Activation-induced CD137 expression on CD4+ T cells transduced with FcRγ alone, Mincle, or Mincle with FcRγ cultured for 18 h with trehalose-6,6-dibehenate (TDB) at the indicated concentrations.

**Supplementary Table 1:**
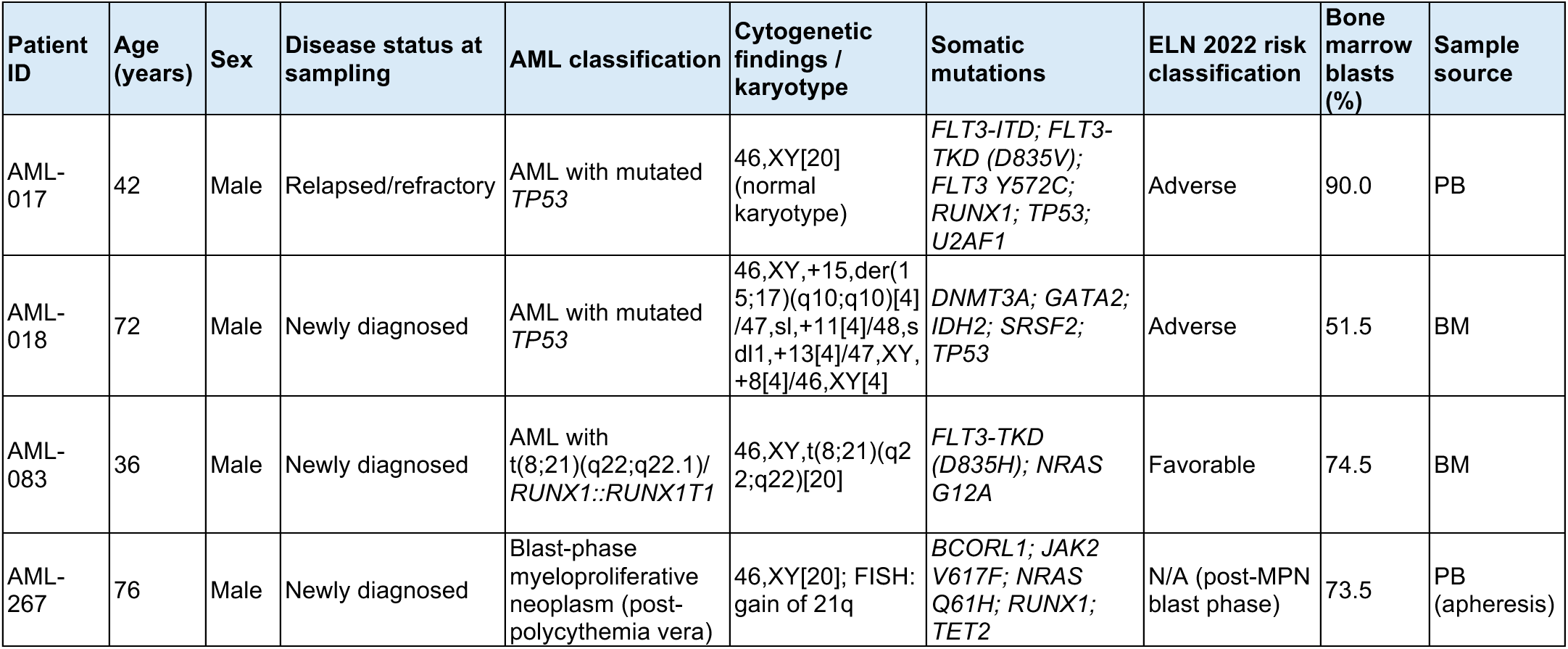
Characteristics of patients with AML and primary samples used in this study. BM, bone marrow; PB, peripheral blood; ELN, European LeukemiaNet. AML classification terminology follows the 2022 ELN/International Consensus Classification^52^. Pathogenic TP53 variants with variant allele frequency >=10% take precedence over myelodysplasia-related gene or cytogenetic categories. AML-267 arose from JAK2-mutated polycythemia vera; per ELN 2022, cases with a prior MPN are classified as accelerated/blast-phase MPN rather than AML. Only pathogenic/likely pathogenic molecular alterations are listed; variants of uncertain significance are omitted. Patient identifiers are study-specific and de-identified.

## References

1 Cappell, K. M. & Kochenderfer, J. N. Long-term outcomes following CAR T cell therapy: what we know so far. Nature Reviews Clinical Oncology 20, 359–371 (2023). 10.1038/s41571-023-00754-1

2 Patel, K. K., Tariveranmoshabad, M., Kadu, S., Shobaki, N. & June, C. From concept to cure: The evolution of CAR-T cell therapy. Molecular Therapy 33, 2123–2140 (2025). 10.1016/j.ymthe.2025.03.005

3 Ruf, B., Greten, T. F. & Korangy, F. Innate lymphoid cells and innate-like T cells in cancer - at the crossroads of innate and adaptive immunity. Nat Rev Cancer 23, 351–371 (2023). 10.1038/s41568-023-00562-w

4 Aguilar, O. A., Fong, L. K. & Lanier, L. L. ITAM-based receptors in natural killer cells. Immunol Rev 323, 40–53 (2024). 10.1111/imr.13313

5 Zhang, Y., Hu, R., Xie, X. & Li, Y. Expanding the frontier of CAR therapy: comparative insights into CAR-T, CAR-NK, CAR-M, and CAR-DC approaches. Ann Hematol 104, 4305–4317 (2025). 10.1007/s00277-025-06538-0

6 Lei, W. et al. Safety and feasibility of 4-1BB co-stimulated CD19-specific CAR-NK cell therapy in refractory/relapsed large B cell lymphoma: a phase 1 trial. Nat Cancer 6, 786–800 (2025). 10.1038/s43018-025-00940-3

7 Lanier, L. L., Yu, G. & Phillips, J. H. Analysis of Fc gamma RIII (CD16) membrane expression and association with CD3 zeta and Fc epsilon RI-gamma by site-directed mutation. J Immunol 146, 1571–1576 (1991).

8 Lanier, L. L., Corliss, B. C., Wu, J., Leong, C. & Phillips, J. H. Immunoreceptor DAP12 bearing a tyrosine-based activation motif is involved in activating NK cells. Nature 391, 703–707 (1998). 10.1038/35642

9 Pende, D. et al. Identification and molecular characterization of NKp30, a novel triggering receptor involved in natural cytotoxicity mediated by human natural killer cells. J Exp Med 190, 1505–1516 (1999). 10.1084/jem.190.10.1505

10 Robinson, M. J. et al. Dectin-2 is a Syk-coupled pattern recognition receptor crucial for Th17 responses to fungal infection. J Exp Med 206, 2037–2051 (2009). 10.1084/jem.20082818

11 Clémenceau, B. et al. Antibody-dependent cellular cytotoxicity (ADCC) is mediated by genetically modified antigen-specific human T lymphocytes. Blood 107, 4669–4677 (2006). 10.1182/blood-2005-09-3775

12 Zhang, T., Wu, M. R. & Sentman, C. L. An NKp30-based chimeric antigen receptor promotes T cell effector functions and antitumor efficacy in vivo. J Immunol 189, 2290–2299 (2012). 10.4049/jimmunol.1103495

13 Tal, Y. et al. An NCR1-based chimeric receptor endows T-cells with multiple anti-tumor specificities. Oncotarget 5, 10949–10958 (2014). 10.18632/oncotarget.1919

14 Ollier, J., Vivien, R., Vié, H. & Clémenceau, B. Transfection of FcγRIIIa (CD16) Alone Can Be Sufficient To Enable Human αβTCR T Lymphocytes To Mediate Antibody-Dependent Cellular Cytotoxicity. ImmunoHorizons 1, 63–70 (2017). 10.4049/immunohorizons.1700022

15 Chou, C. et al. Programme of self-reactive innate-like T cell-mediated cancer immunity. Nature 605, 139–145 (2022). 10.1038/s41586-022-04632-1

16 Correia, M. P. et al. Distinct human circulating NKp30(+)FcεRIγ(+)CD8(+) T cell population exhibiting high natural killer-like antitumor potential. Proc Natl Acad Sci U S A 115, E5980–e5989 (2018). 10.1073/pnas.1720564115

17 Delpoux, A., Lai, C.-Y., Hedrick, S. M. & Doedens, A. L. FOXO1 opposition of CD8+ T cell effector programming confers early memory properties and phenotypic diversity. Proceedings of the National Academy of Sciences 114, E8865–E8874 (2017). 10.1073/pnas.1618916114

18 Miron, M. et al. Human Lymph Nodes Maintain TCF-1(hi) Memory T Cells with High Functional Potential and Clonal Diversity throughout Life. J Immunol 201, 2132–2140 (2018). 10.4049/jimmunol.1800716

19 Savage, A. K. et al. The transcription factor PLZF directs the effector program of the NKT cell lineage. Immunity 29, 391–403 (2008). 10.1016/j.immuni.2008.07.011

20 Kovalovsky, D. et al. The BTB-zinc finger transcriptional regulator PLZF controls the development of invariant natural killer T cell effector functions. Nat Immunol 9, 1055–1064 (2008). 10.1038/ni.1641

21 Constantinides, M. G., McDonald, B. D., Verhoef, P. A. & Bendelac, A. A committed precursor to innate lymphoid cells. Nature 508, 397–401 (2014). 10.1038/nature13047

22 Brandt, C. S. et al. The B7 family member B7-H6 is a tumor cell ligand for the activating natural killer cell receptor NKp30 in humans. J Exp Med 206, 1495–1503 (2009). 10.1084/jem.20090681

23 Textor, S. et al. The proto-oncogene Myc drives expression of the NK cell-activating NKp30 ligand B7-H6 in tumor cells. Oncoimmunology 5, e1116674 (2016). 10.1080/2162402x.2015.1116674

24 Chen, H. et al. Deficiency of N-linked glycosylation impairs immune function of B7-H6. Front Immunol 14, 1255667 (2023). 10.3389/fimmu.2023.1255667

25 Miller, J. S. et al. Expansion and homing of adoptively transferred human natural killer cells in immunodeficient mice varies with product preparation and in vivo cytokine administration: implications for clinical therapy. Biol Blood Marrow Transplant 20, 1252–1257 (2014). 10.1016/j.bbmt.2014.05.004

26 Cooper, M. A. et al. In vivo evidence for a dependence on interleukin 15 for survival of natural killer cells. Blood 100, 3633–3638 (2002). 10.1182/blood-2001-12-0293

27 Haubner, S., Subklewe, M. & Sadelain, M. Honing CAR T cells to tackle acute myeloid leukemia. Blood 145, 1113–1125 (2025). 10.1182/blood.2024024063

28 Khalifeh, M., Hopewell, E. & Salman, H. CAR-T cell therapy for treatment of acute myeloid leukemia, advances and outcomes. Mol Ther 33, 2441–2453 (2025). 10.1016/j.ymthe.2025.03.052

29 Letourneur, F. & Klausner, R. D. T-cell and basophil activation through the cytoplasmic tail of T-cell-receptor zeta family proteins. Proc Natl Acad Sci U S A 88, 8905–8909 (1991). 10.1073/pnas.88.20.8905

30 O’Shea, J. J., Weissman, A. M., Kennedy, I. C. & Ortaldo, J. R. Engagement of the natural killer cell IgG Fc receptor results in tyrosine phosphorylation of the zeta chain. Proc Natl Acad Sci U S A 88, 350–354 (1991). 10.1073/pnas.88.2.350

31 Robbins, P. F. et al. Single and dual amino acid substitutions in TCR CDRs can enhance antigen-specific T cell functions. J Immunol 180, 6116–6131 (2008). 10.4049/jimmunol.180.9.6116

32 Pessino, A. et al. Molecular Cloning of NKp46: A Novel Member of the Immunoglobulin Superfamily Involved in Triggering of Natural Cytotoxicity. Journal of Experimental Medicine 188, 953–960 (1998). 10.1084/jem.188.5.953

33 Sen Santara, S., et al. The NK cell receptor NKp46 recognizes ecto-calreticulin on ER-stressed cells. Nature 616, 348–356 (2023). 10.1038/s41586-023-05912-0

34 Lanier, L. L., Yu, G. & Phillips, J. H. Co-association of CD3 zeta with a receptor (CD16) for IgG Fc on human natural killer cells. Nature 342, 803–805 (1989). 10.1038/342803a0

35 Kraft, S. & Kinet, J.-P. New developments in FcεRI regulation, function and inhibition. Nature Reviews Immunology 7, 365–378 (2007). 10.1038/nri2072

36 Deng, M., Du, S., Hou, H. & Xiao, J. Structural insights into the high-affinity IgE receptor FcεRI complex. Nature 633, 952–959 (2024). 10.1038/s41586-024-07864-5

37 Ward, D. E., Fay, B. L., Adejuwon, A., Han, H. & Ma, Z. Chimeric Antigen Receptors Based on Low Affinity Mutants of FcεRI Re-direct T Cell Specificity to Cells Expressing Membrane IgE. Frontiers in Immunology **Volume** 9–2018 (2018). 10.3389/fimmu.2018.02231

38 Yonekawa, A. et al. Dectin-2 is a direct receptor for mannose-capped lipoarabinomannan of mycobacteria. Immunity 41, 402–413 (2014). 10.1016/j.immuni.2014.08.005

39 Sato, K. et al. Dectin-2 is a pattern recognition receptor for fungi that couples with the Fc receptor gamma chain to induce innate immune responses. J Biol Chem 281, 38854–38866 (2006). 10.1074/jbc.m606542200

40 Saijo, S. et al. Dectin-2 recognition of alpha-mannans and induction of Th17 cell differentiation is essential for host defense against Candida albicans. Immunity 32, 681–691 (2010). 10.1016/j.immuni.2010.05.001

41 Ishikawa, T. et al. Identification of distinct ligands for the C-type lectin receptors Mincle and Dectin-2 in the pathogenic fungus Malassezia. Cell Host Microbe 13, 477–488 (2013). 10.1016/j.chom.2013.03.008

42 Schoenen, H. et al. Cutting edge: Mincle is essential for recognition and adjuvanticity of the mycobacterial cord factor and its synthetic analog trehalose-dibehenate. J Immunol 184, 2756–2760 (2010). 10.4049/jimmunol.0904013

43 Ishikawa, E. et al. Direct recognition of the mycobacterial glycolipid, trehalose dimycolate, by C-type lectin Mincle. J Exp Med 206, 2879–2888 (2009). 10.1084/jem.20091750

44 Yamasaki, S. et al. Mincle is an ITAM-coupled activating receptor that senses damaged cells. Nat Immunol 9, 1179–1188 (2008). 10.1038/ni.1651

45 Irving, B. A., Chan, A. C. & Weiss, A. Functional characterization of a signal transducing motif present in the T cell antigen receptor zeta chain. J Exp Med 177, 1093–1103 (1993). 10.1084/jem.177.4.1093

46 Johnson, S. A. et al. Phosphorylated immunoreceptor signaling motifs (ITAMs) exhibit unique abilities to bind and activate Lyn and Syk tyrosine kinases. J Immunol 155, 4596–4603 (1995).

47 Koene, H. R. et al. FcγRIIIa-158V/F Polymorphism Influences the Binding of IgG by Natural Killer Cell FcγRIIIa, Independently of the FcγRIIIa-48L/R/H Phenotype. Blood 90, 1109–1114 (1997). 10.1182/blood.V90.3.1109

48 Jutz, S. et al. Assessment of costimulation and coinhibition in a triple parameter T cell reporter line: Simultaneous measurement of NF-κB, NFAT and AP-1. J Immunol Methods 430, 10–20 (2016). 10.1016/j.jim.2016.01.007

49 Seet, C. S. et al. Generation of mature T cells from human hematopoietic stem and progenitor cells in artificial thymic organoids. Nat Methods 14, 521–530 (2017). 10.1038/nmeth.4237

50 Han, J., Tam, K., Tam, C., Hollis, R. P. & Kohn, D. B. Improved lentiviral vector titers from a multi-gene knockout packaging line. Mol Ther Oncolytics 23, 582–592 (2021). 10.1016/j.omto.2021.11.012

51 Lanigan, T. M. et al. Real time visualization of cancer cell death, survival and proliferation using fluorochrome-transfected cells in an IncuCyte(®) imaging system. J Biol Methods 7, e133 (2020). 10.14440/jbm.2020.323

52 Döhner, H. et al. Diagnosis and management of AML in adults: 2022 recommendations from an international expert panel on behalf of the ELN. Blood 140, 1345–1377 (2022). 10.1182/blood.2022016867

